# High-resolution metabolite imaging: luteolinidin accumulates at the host-cyanobiont interface during cold-acclimation in *Azolla* symbioses

**DOI:** 10.1101/2024.01.05.574342

**Authors:** Erbil Güngör, Benjamin Bartels, Giorgio Bolchi, Ron M.A. Heeren, Shane R. Ellis, Henriette Schluepmann

**Author notes:** Current address for Benjamin Bartels: Biomolecular Chemistry, Leibniz Institute for Natural Product Research and Infection Biology / Hans Knöll Institute, Adolf-Reichwein-Straße 23, 07745 Jena, Germany.

## Abstract

Aquatic ferns of the genus *Azolla* (Azolla) form highly productive symbioses with filamentous cyanobacteria fixing N_2_ in their leaf cavities, *Nostoc azollae*. Stressed symbioses characteristically turn red due to 3-deoxyanthocyanin (DA) accumulation, rare in angiosperms and of unknown function. To reveal DA functions upon cold acclimation and recovery, we integrated laser-desorption-ionization mass-spectrometry-imaging (LDI-MSI), a new *A. filiculoides* genome-assembly and annotation, and dual RNA-sequencing into phenotypic analyses of the symbioses.

*Azolla sp.* Anzali recovered even when cold-induced DA-accumulation was inhibited by abscisic acid. Cyanobacterial filaments generally disappeared upon cold acclimation, and *N. azollae* transcript profiles were unlike those of resting stages formed in cold-resistant sporocarps, yet filaments re-appeared in leaf cavities of newly formed green fronds upon cold-recovery.

The high transcript accumulation upon cold acclimation of *AfDFR1* encoding a flavanone 4-reductase active *in vitro* suggested that the enzyme of the first step in the DA-pathway may regulate accumulation of DAs in different tissues. However, LDI-MSI highlighted the necessity to describe metabolite accumulation beyond class assignments as individual DA and caffeoylquinic acid metabolites accumulated differentially. For example, luteolinidin accumulated in epithelial cells, including those lining the leaf cavity, supporting a role for the former in the symbiotic interaction during cold acclimation.

**Summary statement:** During cold acclimation in *Azolla* symbioses, individual compounds from the same phenolic class accumulated in different host tissues: luteolinidin associated with biotic interactions at the symbiosis interface whilst apigenidin with photooxidative stress mitigation in the mesophyll.

## Introduction

The prolifically growing Azolla fern symbioses do not require any nitrogen fertilizer and their biomass has proven suitable for inclusion in feed formulations, therefore, they are currently considered for protein cropping in delta regions throughout the world (Brouwer et al., 2018). The high content of phenolics in their biosmass, particularly protein-binding proanthocyanidins (PAs), however, limits its rate of inclusion into feed formulations. In fact, phenolics effectively protect from grazers and likely are of importance for symbiont-host interactions (Brouwer et al., 2019; Güngör et al., 2023). Currently, our fundamental knowledge about the role individual phenolic compounds play in symbiotic and defense interactions under different growth conditions is not sufficient to guide in the domestication of Azolla.

It is known, however, that phenolics crucially affect plant-microbe interactions in seed plants, particularly so the class of flavonoids, directly or after modification by microbes. Plant-microbe symbioses are no exceptions, as shown in the researched cases of rhizobia or arbuscular mycorrhizal fungi with seed plants (Wang et al., 2022). Cyanobacterial symbioses with plants have not been studied to the same extent. In addition, Azolla symbioses differ from the root symbioses studied thus far in that the cyanobacterial colony is maintained at the shoot apex and inoculates leaves and sporocarps during their early development (Dijkhuizen et al., 2021; Güngör et al., 2023). Inside the fully matured leaf cavities, *N. azollae* filaments frequently form heterocysts and fix sufficient N_2_ for even the highest growth rates of the symbiosis (Brouwer et al., 2017). *Azolla filiculoides* thrives in temperate climates, is common in the Netherlands and was chosen for genome sequencing and annotation thus providing knowledge required for molecular studies in 2018 (Pieterse et al., 1977; Li et al., 2018). It is winter-hardy (**Figure 1A**), and turns completely red in cold due to the accumulation of red flavonoid 3-deoxyanthocyanins (DAs) (Iwashina et al., 2010). DAs are generally attributed to ferns (Harborne, 1965; Piatkowski et al., 2020), but also occur in isolated species of mono– and dicotyledonous angiosperms, such as *Sorghum bicolor* and *Sinningia cardinalis*, respectively (Awika et al., 2004; Winefield et al., 2005). DAs differ from the well-studied 3-hydroxyanthocyanins (HAs) by the lack of one hydroxyl-group at position 3 of the C-ring (**Figure 1B**). DAs therefore do not show radical color changes at pH extremes and are more resistant to high temperatures compared to HAs (Alejo-Armijo et al., 2020; Herrman et al., 2020). In spite of their qualities, DA biosynthesis, (sub)cellular localization and the function of specific DAs remains unknown.

**Figure 1.**
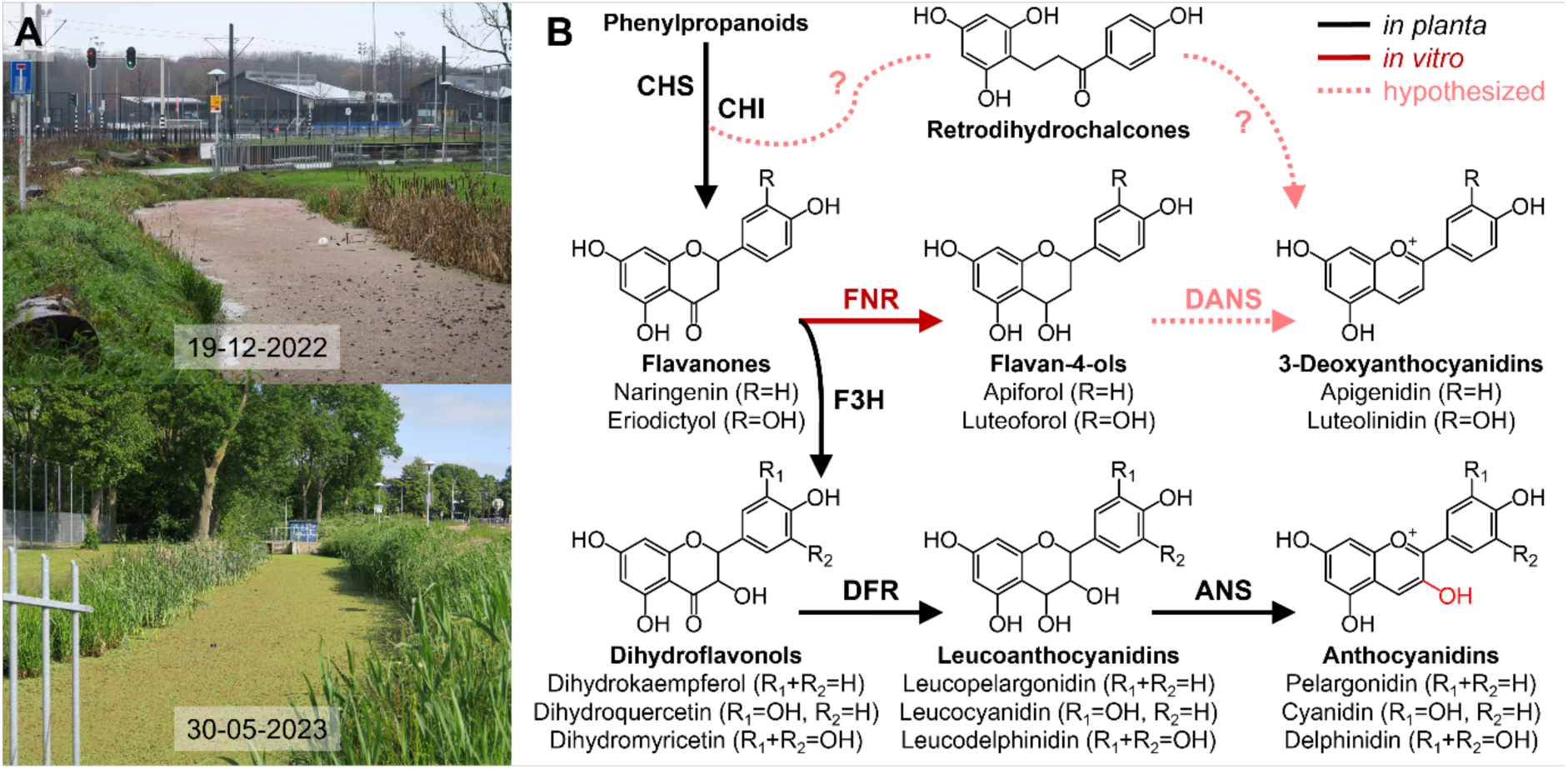
Characteristic accumulation of red 3-deoxyanthocyanidins and derived glucosides (DAs) in Azolla. **A** *A. filiculoides* in the Galgenwaard ditch (Utrecht, The Netherlands) in winter (19-12-2022) and in spring (30-05-2023) provided the strain of which the genome was sequenced. **B** Proposed pathways for DA-production. Analogous to angiosperm anthocyanidin biosynthesis, DA may be synthesized with two enzymes starting from flavanones: (1) flavanone 4-reductase (FNR) and (2) deoxyanthocyanidin synthase (DANS). FNR-activity for a dihydroflavonol 4-reductase (DFR) homolog from *Sorghum bicolor* was verified *in vitro* (Kawahigashi et al., 2016), but the second enzyme has not yet been found. Alternatively, a pathway via retrodihydrochalcone intermediates has been proposed (Khalil et al., 2012). Enzymes, CHS: chalcone synthase, CHI: chalcone isomerase, F3H: flavanone 3-hydroxylase, and ANS: anthocyanidin synthase.

The main functions attributed to stress-induced red flavonoid pigment accumulation in land plants are photoprotection by filtering blue/green light, and ROS-scavenging (Davies et al., 2022). Consistently, excess light triggered DA-accumulation in *A. filiculoides* (Brilli et al., 2022), often in combination with abiotic stresses such as nutrient deficiency or cold (Tran et al., 2020; Costarelli et al., 2021). Prolonged cold-exposure may distort the carbon and nitrogen cycling within the symbioses and cause photooxidative damage that may result in starvation of the host. As a further implication, overwintering *N. azollae* symbionts may differentiate into the cold-resistant single-celled resting stages, akinetes, storing nitrogenous compounds in the form of cyanophycin (Garg & Maldener, 2021). Akinetes of *Nostoc sp.* PCC 7524 survived for 15 months at 4°C, whilst the vegetative cells died after a week (Sutherland et al., 1979). DAs extracted from Azolla have indeed been reported to affect cellular differentiation in a free-living Nostoc model with altered expression of motility genes in *Nostoc punctiforme* (Cohen et al., 2002a). Molecular processes in *N. azollae* when symbioses acclimate to the cold have not been described.

Key flavonoid biosynthesis pathway (FBP) enzymes for HA-production are dihydroflavonol reductase (DFR) and anthocyanidin synthase (ANS) (Wilmouth et al., 2002; Petit et al., 2007). A homolog of DFR, flavanone 4-reductase (FNR), has been proposed as the first enzyme in DA-biosynthesis from flavanones (**Figure 1B**). An FNR was upregulated when DAs accumulated in *S. bicolor,* and converted flavanones into flavan-4-ols, which was converted into DAs by acid-heat treatment *in vitro* (Halbwirth et al., 2003; Kawahigashi et al., 2016). Since both leucoanthocyanidins and flavan-4-ols produced by DFR and FNR, respectively, turn into HAs and DAs by acid-heat treatment, the second catalytic step in DA-production was also attributed to ANS-activity (Winefield et al., 2005). The two-enzyme hypothesis is further supported by a similar regulation of the transcription of the enzymes: the same environmental stress conditions that induce DA-production in Azolla, induce HA-production in *Arabidopsis thaliana* via R2R3-MYB transcription factor complexes (Schulz et al., 2016). Yet an ANS enzyme carrying out the second conversion in DA biosynthesis has never been found in Sorghum. In addition, the hydroxyl-group at position C3, lacking in flavan-4-ols, is considered essential for ANS-activity (Wilmouth et al., 2002) and ANS was downregulated during DA-accumulation in *S. bicolor* (Hipskind, 1996; Liu et al., 2010). Alternatively, a retrodihydrochalcone has been proposed as the substrate for DA-production, as it was detected in *S. bicolor* extracts and behaved the same as the flavan-4-ol apiforol during acid-heat treatment (Khalil et al., 2012).

In *S. bicolor,* DAs are considered key phytoalexins as they are rapidly produced locally in response to cell damage (Stutts & Vermerris, 2020). *S. bicolor yellow seed1* mutants lacking DAs showed higher susceptibility to corn leaf aphid *Rhopalosiphum maidis* and the anthracnose fungus *Colletotrichum sublineolum* (Ibraheem et al., 2010; Kariyat et al., 2019). Azolla DAs might therefore also mediate chemical defense. Similarly, *Lymnea swinhoei* snails and *Polypedates leucomystax* tadpoles preferred feeding on *A. filiculoides* fronds with low DA-concentration over *A. pinnata* fronds with high DA-concentration (Cohen et al., 2002b). The preference could, however, also be due to *A. pinnata* in general having a higher PA-concentration than *A. filiculoides* (Brouwer et al., 2019; Güngör et al., 2021). Considering the small size of Azolla, it would be too late if phytoalexins were only produced after cell damage has occurred. Azolla, therefore, presumably, produces high concentrations of phenolics prophylactically (Brouwer et al., 2018). In this way Azolla usually manages to outgrow its enemies with the exception of the weevil *Stenopelmus rufinasus* (Hill, 1998). *S. rufinasus* larvae have been reported to turn red when feeding on red *A. filiculoides* (Richerson & Grigarick, 1967).

It remains challenging to pinpoint which phenolic does what, as phenolics often share functional groups, and are induced hand-in-hand during environmental stress(es) (Hadacek et al., 2011). Azolla also produces high concentrations of caffeoylquinic acids (CQAs), which have comparable roles to PAs and DAs (Qian et al., 2020; Alcázar Magaña et al., 2021). CQAs are a product of the core phenylpropanoid biosynthesis pathway (PBP) shared by CQAs, PAs and DAs. Therefore, investment of aromatic acids into the different types of phenolics is very likely finely regulated in the fern host. Conventional analyses of whole plant extracts have often overlooked localization. Mapping the spatial distribution of molecules with laser-desorption-ionization mass-spectrometry-imaging (LDI-MSI) could therefore tell us more about the possible functions of phenolics in the Azolla symbiosis (McDonnell & Heeren, 2007; Boughton et al., 2016).

Here we investigate, how temperate and (sub)tropical Azolla symbioses respond to and recover from different cold treatments, and simultaneously follow the response of the endosymbiont *N. azollae*. We then map the spatial distribution of accumulating phenolics, comparing cold-treated and green *A. filiculoides* with LDI-MSI. Leveraging a new *A. filiculoides* genome assembly and annotation (Afi_v2), we further analyze the gene expression of *A. filiculoides* and *N. azollae* in response to cold with dual RNA-sequencing. We finish by characterizing possible enzymes of DA-biosynthesis using a phylogenomic analysis of the 2-oxoglutarate dependent dioxygenase superfamily, recombinant enzyme assays and *in planta* complementation.

## Materials and methods

### Plant materials

The four Azolla species used in this study were *A. filiculoides* strain Galgenwaard (Li et al., 2018), *A. pinnata* originating from the Botanical Gardens of Antwerp (Belgium), an *Azolla* species from the Anzali lagoon (Iran), which could not be phylogenetically assigned to any of the described Azolla species (Dijkhuizen et al., 2021), and an unknown Azolla species collected from the Botanical Gardens of Bordeaux (France).

### Growth conditions and cold-treatment

Standard growth medium contained 0.65 mM NaH_2_PO_4_·H_2_O, 1.02 mM K_2_SO_4_, 1 mM CaCl_2_·2H2O, 1.65 mM MgSO_4_·7H_2_O, 17.9 µM Fe-EDTA, 9.1 µM MnCl_2_·4H_2_O, 1.6 µM Na_2_MoO_4_·2H_2_O, 18.4 µM H_3_BO_3_, 0.8 µM ZnSO_4_·7H_2_O, 0.3 µM CuSO_4_·5H_2_O and 0.2 µM CoCl_2_·6H_2_O. Variations did not contain NaH_2_PO_4_·H_2_O or were supplemented with 5 µM abscisic acid. Optimal growth conditions were at 21°C, 70% humidity and 16 h light (or 12 h if specified) with an intensity of 100 µmol m^−2^ s^−1^, resulting in green sporophytes. Cold-treatment was at 4°C, 70% humidity and 16 h (or 12 h if specified) light with an intensity of 100 µmol m^−2^ s^−1^, resulting in red sporophytes.

### Estimation of 3-deoxyanthocyanin content

Fresh *A. filiculoides* sporophytes (50 mg) were snap-frozen and mechanically sheared for 50 sec (30 Hz). The ground tissue was extracted for 1.5 h at 4°C with 1.2 ml methanol-HCl (0.5% v/v). The suspension was centrifuged for 5 min at 10 000 g, and the supernatant was diluted 1:1 with methanol before the absorbance was measured at 500 nm. Extracts had a peak absorbance at 490 nm, close to that of luteolinidin at 500 nm. Luteolinidin (TransMIT, Giessen, Germany) was therefore used as standard to estimate the DA-content in µg per g FW.

### Microscopy

Images of Azolla shoot apices, anthocyanic vacuolar inclusions, *N. azollae* and leaf pockets were taken with a Zeiss Axio Zoom.V16 microscope (Zeiss, Jena, Germany) with a Zeiss Axiocam 506 color camera, Zeiss CL 9000 LED lights and a Zeiss HXP 200C fluorescence lamp with standard Zeiss RFP filter set 63HE (excitation 572 nm, emission 629 nm). Individual *N. azollae* filaments were exposed by crushing Azolla shoot apices between a cover slip and a microscope slide with a drop of demineralized water. Images were Z-stacked with Helicon Focus 8 software in default settings (depth map; radius 8, smoothing 4).

### Leaf pocket isolation

The methods was as in Güngör et al., 2023. Briefly, sporophytes were treated with cellulase, macerozyme and pectolyase for 19 h at 30°C after which leaf pockets were mechanically released and manually collected. Sporophytes were only cold-treated for one week before isolation, because after a longer treatment oxidative browning during enzymatic digestion became unavoidable.

### Visualization of proanthocyanidins

A method was adapted from Abeynayake et al., 2011. Briefly, sporophytes were decolorized for 2 h in absolute ethanol and stained for 20 min with 0.01% (w/v) 4-dimethylaminocinnamaldehyde (DMACA) dissolved in ethanol-HCl (0.5% v/v). Stained sporophytes turned turquoise and were fixated for 2 h at 4°C with 6% (w/v) glutaraldehyde and 4% (w/v) paraformaldehyde dissolved in 50 mM sodium phosphate buffer (pH 7.4). Sporophytes turned purple after fixation and were immersed in absolute ethanol until imaging.

### Sample preparation for mass spectrometry imaging

*A. filiculoides* sporophytes were either grown in optimal growth conditions, or cold-treated for three weeks. Before embedding, plants were submerged in demineralized water with 0.01% (v/v) Silwet L-77 (Lehle Seeds, Gent, Belgium) until they sunk underneath the water surface (5-10 sec). Excess solution was absorbed with a paper towel. Sporophytes from both conditions were placed together in a mold, and embedded in 20% (w/v) gelatin. The gelatin was supplemented gradually, to trap the plants in the center and minimize air bubble formation. The samples were frozen over dry ice. A Microm HM525 cryostat (Thermo Fisher, Waltham, MA, USA) was used for making 16 µm thick sections, at –20°C. Sections were collected with double-sided adhesive tape inspired by Kawamoto, 2003, and adhered to SuperFrost Plus microscope slides (VWR, Amsterdam, Netherlands), which had been cleaned by sonication, immersed in hexane and ethanol, sequentially. The mounted sections were stored at –80°C until measurement.

### Laser-desorption-ionization mass-spectrometry-imaging (LDI-MSI)

The samples were thawed to room temperature in a desiccator individually before analysis and a 20x optical image was acquired with an Aperio CS2 slide scanner (Leica, Wetzlar, Germany) prior to the MSI experiment. LDI-MSI measurements were performed with a MALDI/ESI injector ion source (Spectroglyph, Kennewick, WA, USA) coupled to an Orbitrap Elite mass spectrometer (Thermo Fisher, Bremen, Germany). Mass spectra were acquired in positive ion mode at a pixel size of 20 µm, and a mass resolving power of 120 000 @ m/z 400. The laser for desorption and ionization sampled each position within the region of interest with 100 pulses and operated at wavelength 349 nm, a fluency of ∼2.7 J/cm^2^ and a repetition rate of 1 kHz (pulse energy 1-3 uJ and spot size 1-3 micron).

A pattern file required for visualizing spatial distributions was produced by Spectroglyph software during acquisition. Raw files with acquired mass spectra data were converted to mzML format with MSConvert 3 software (Chambers et al., 2012) and subsequently to imzML format with LipostarMSI (Tortorella et al., 2020). The latter was used to calculate the spatial distribution of all the detected *m/z* features, applying quantile thresholding (high quantile 99.75) for hotspot removal, and normalization to total ion count, per pixel. Putative chemical identities were assigned to *m/z* features based on accurate mass and isotopic pattern, making use of previously published literature about mass spectrometry experiments on Azolla species (Güngör et al., 2021). Further validation of selected compounds, namely apigenidin, luteolinidin, aesculin, catechin, epicatechin, procyanidin A1 and procyanidin B2, was based on the analysis of commercially available standards of the same compounds (TransMIT) under comparable experimental conditions. For this, all standards were dissolved in Methanol and sprayed onto an ITO glass slide using a TM-sprayer (HTX Technologies, LLC, Chapel Hill NC, USA) and then subjected to an LDI-MSI experiment.

### Dual RNA-sequencing and data analysis

Green *A. filiculoides* sporophytes were grown in 500 ml standard medium at 4°C, with a shortened light period of 12 h to have a more gradual course of DA-production, for either 14 days or 21 days (cold-treated day 14, **C14**; cold-treated day 21, **C21**). Control sporophytes were grown in parallel at 21°C for 14 days (untreated day 14, **U14**). All conditions were sampled in triplicate, 2 h into the light cycle of the diel rhythm. Dual RNA-Seq data from the spores (sporophyte, **sp**; megasporocarp, **mega**; microsporocarp, **micro**) was already available as described in Dijkhuizen et al., 2021 and Güngör et al., 2023.

Total RNA was isolated with the Spectrum Plant Total RNA Kit (Sigma-Aldrich, Saint-Louis, MO, USA) using protocol B and treated with DNase I (Thermo Fisher) for 1 h at 37°C after which the reaction was stopped by 10 min. 65°C treatment with EDTA and cleaned with the RNeasy MinElute Cleanup Kit (Qiagen, Hilden, Germany). Depletion of ribosomal RNA (rRNA) was carried out per instructions of a custom designed riboPOOL Kit (siTOOLs-Biotech, Planegg, Germany) for *A. filiculoides* and *N. azollae* rRNA and cleaned again with the aforementioned kit. Stranded libraries were synthesized using the Universal Prokaryotic RNA-Seq AnyDeplete Kit (Tecan, Männedorf, Switzerland), quality checked with TapeStation DNA ScreenTape (Agilent Technologies, USA) and then sequenced on half lane NovaSeq 6000 using the paired-end 2×50 cycle chemistry (Illumina, San Diego, CA, USA). Data is deposited under accession number PRJEB60373.

Acquired reads were demultiplexed, quality-filtered and trimmed to remove primers before alignment with STAR to the concatenated genomes of *A. filiculoides* v2, its chloroplast and *N. azollae* (Dobin et al., 2013; Güngör et al., 2023). For the spore samples only reads for *N. azollae* were extracted whilst for the cold-treatment reads were extracted for both *N. azollae* and *A. filiculoides*. Based on PCA-plots (**File S1**), outlier samples **C21-3**, **micro-2** and **mega-3** were removed for *N. azollae*, before calculating differential gene expression with DESeq2 in default settings with adaptive shrinkage to remove low expressors with high dispersion values (Love et al., 2014). Replicates of the cold-treated *A. filiculoides* did not have outliers and were all used for statistical analysis with DESeq2 (**File S2**).

### Cloning of *Af*DFR1

We used Golden Gate Cloning (Engler et al., 2014) for cloning *Af*DFR1. The gene was PCR-amplified from *A. filiculoides* cDNA (Fragment 1 F-primer: ATGAAGACATAATGGCAATTACTGACTCCGCCAA, R-primer: ATGAAGACATTTTTCTTTAAATCATTCGCATCAC; Fragment 2 F-primer: ATGAAGACATAAAACGGCACTTCTAAGAGAACTC, R-primer, ATGAAGACATAAGCTCATGGGGTACTTCCAGGAT) in two fragments to eliminate an internal restriction site by silent mutation of 183G>A and domesticated into level 0. CaMV35S-promotor, N-terminal 6xHis-tag and AtuNos-terminator were cloned together with the gene in level 1, and subsequently cloned into the final level 2 (pAGM4673) including green fluorescent protein (GFP) with the same promotor/terminator and a readily available kanamycin-resistance cassette (pICSL70004). The final plasmid was transformed into *Agrobacterium tumefaciens* strain GV3101 through electroporation.

### Procedures for *in vitro* enzyme assay

For assaying substrate-specificity of *Af*DFR1, the protein was recombinantly expressed in *Nicotiana tabacum* leaves through leaf-infiltration with *Agrobacterium tumefaciens* (Sheludko et al., 2007). Briefly, a solution containing *A. tumefaciens*, 1% sucrose, 0.005% Silwet L-77, ¼MS and 100 µM acetosyringone was injected into adult *Nicotiana tabacum* leaves with a syringe. GFP-expression was visible after 72 h, and as both proteins were driven by the same promotor, we used this time point for isolating the enzyme.

All procedures from hereon were performed at 4°C, if possible. Transformed leaves were cut in small pieces and ground in pestle mortar with 10 ml extraction buffer per 1 gram of material (0.1 M borate pH 8.8, 0.4 M NaCl, 10% glycerol (v/v), 2.5% polyvinylpyrrolidone (w/v), 20 mM sodium ascorbate, 1 mM PMSF and 1 mM TCEP) and centrifuged at 4700 g for 30 min before loading the supernatant on a HisPur Ni-NTA Spin Column (Thermo Fisher). The column with cleared lysate was rotated for 1 h after which it was washed three times with washing buffer (50 mM Tris-HCl pH 7.5, 500 mM NaCl, 10 mM imidazole, 5% glycerol). Enzyme reaction volume was 1 ml and contained 50 mM Tris-HCl pH 7.5, 1 mM substrate (naringenin, eriodictyol or dihydroquercetin, TransMIT) and 1 mM NADPH (Sigma-Aldrich), and was added directly to the column, incubated for 1 h at 37°C and centrifuged. Hereafter, the enzyme reactions were stopped by extracting twice with 500 µl ethyl acetate and vacuum-evaporated. Acid-heat treatment was performed by adding 50 µl methanol-HCl (5% v/v) to the pellet and a subsequent incubation for 5-10 sec at 90°C, after which the red color developed. Samples for LC-MS measurement were split in two before vacuum-evaporation, after which one pellet was dissolved in ultrapure water and the other was acid-heat treated as described above. Samples were stored at –80°C until LC-MS analysis as described by Kawahigashi et al., 2016.

### Procedures for *in planta* complementation

*Arabidopsis thaliana tt3* mutants were requested from NASC (ID:N84) and stably transformed by the floral-dip transformation method (Zhang et al., 2006). Briefly, flowers were dipped for 5-10 sec in a solution containing *A. tumefaciens*, 5% sucrose and 0.01% Silwet L-77 and left to set seed. The harvested seeds were selected on 0.8% plant agar ½MS plates (pH 5.8) with 100 mg/L kanamycin and GFP-expression in leaves of resistant plants was verified. HA-accumulation under high light intensity and the seed color of three individual transgenic plants was used as a measure for HA-production.

## Results

### Cold-treatment in relation to cold-recovery in different Azolla symbioses

Shifting *A. filiculoides* sporophytes from ambient 21°C to 4°C reliably induced red pigmentation: the plants stopped growing and developed a red hue after a week with a 16 h, or after two weeks with a 12 h light period. Supplementing 5 µM abscisic acid (ABA) delayed the appearance of the red hue by a week (**Figure 2A**). Sporulating sporophytes tended to wither after the cold shift and therefore only vegetatively growing ferns were used.

**Figure 2.**
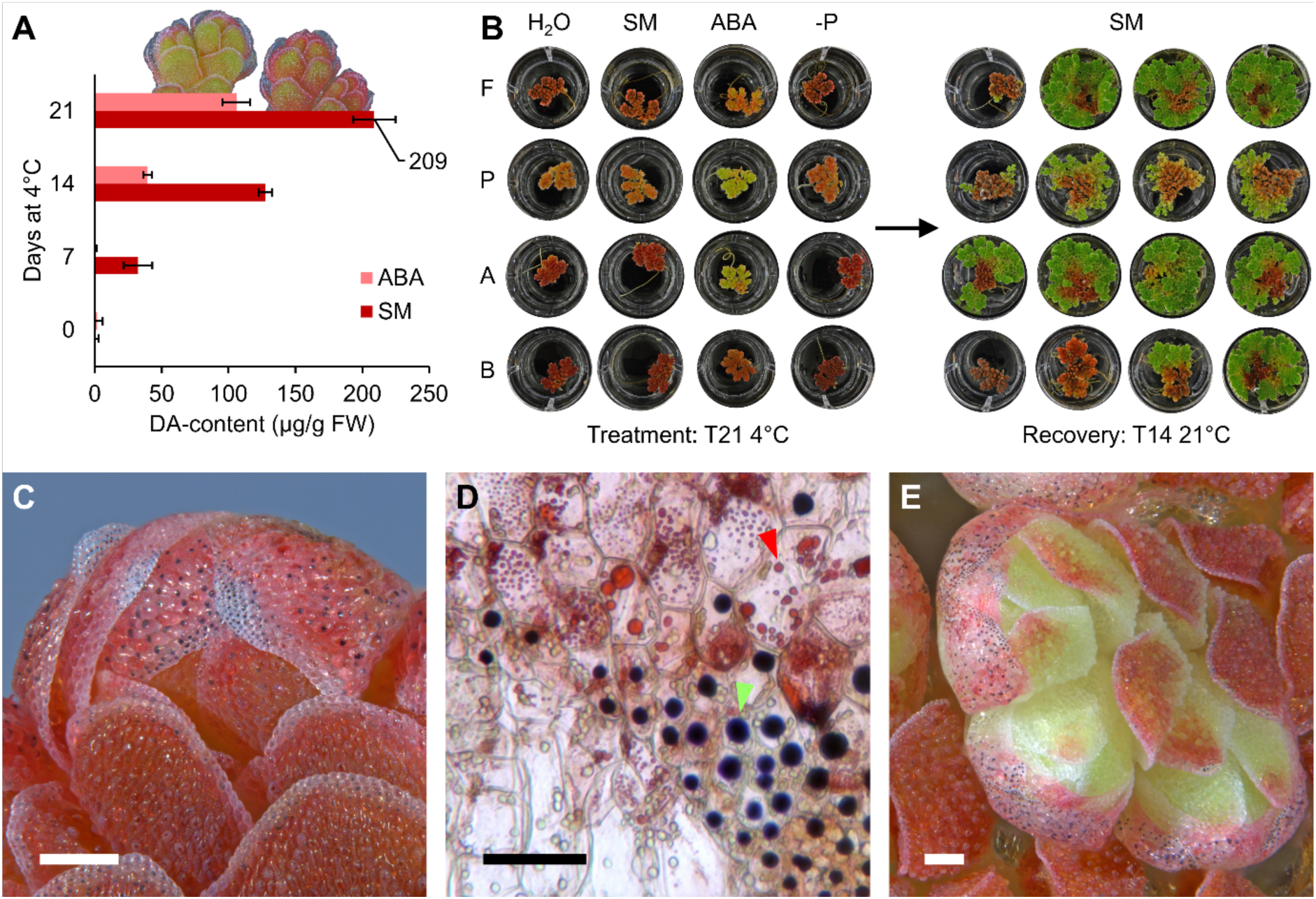
Azolla during cold-treatment at 4°C and recovery at 21°C. **A** Progression of DA-accumulation in *A. filiculoides* on standard medium (SM) without versus with 5 µM abscisic acid (ABA). **B** Typical growth habit of different Azolla species after 21 days 4°C cold-treatment on demineralized water (H_2_O), SM, SM with 5 µM ABA and SM without phosphate (-P), and after 14 days of recovery at 21°C on SM. Species, A: *A. filiculoides*, P: *A. pinnata*, A: *Azolla sp. Anzali*, B: *Azolla sp. Bordeaux*. Full timeline available in **Figure S2**. **C** Typical *A. filiculoides* shoot apex after 21 days 4°C cold-treatment. **D** Close-up of diffused red pigmentation throughout the cells and anthocyanic vacuolar inclusions (AVIs) in different shapes: small, purple (red arrow); big, black (green arrow). **E** Typical increase in AVIs at the leaf rim, after one week recovery at 21°C of a shoot as in **C**. Scale bars: white and black, respectively, correspond to 200 µm and 50 µm.

After 60 days at 4°C, *A. filiculoides* turned completely red for most plants whilst some withered. When moved back to 21°C the redness faded and over the course of two weeks green outgrowth emerged from the shoot apices; after five weeks the whole medium surface was covered again by newly grown green Azolla (**Figure S1**). Since red Azolla tissue was not reviving, we determined recovery success by the time it takes for green growth points to appear.

To compare the temperate *A. filiculoides*, tropical *A. pinnata*, and subtropical *Azolla sp.* Anzali and *Azolla sp.* Bordeaux, we placed plants three weeks at 4°C, then returned to 21°C on fresh standard medium (**Figure 2B, Figure S2**). The medium during cold-treatment varied between demineralized water, SM and SM with ABA or without phosphate. All species on demineralized water turned as red as those on SM, but none except *Azolla sp.* Anzali recovered after two weeks at 21°C (**Figure 2B**). For cold-recovery, DA-content was less important than mineral nutrient availability during the cold. All species on SM without phosphate recovered; *Azolla sp.* Bordeaux was even boosted by a sudden phosphate availability during recovery (**Figure 2B**). Phosphate limitation therefore did not influence cold tolerance. Tropical *A. pinnata* recovered the worst under all conditions, compared to *A. filiculoides* or *Azolla sp.* Anzali (**Figure 2B**). All species accumulated the least DAs with ABA (**Figure 2B**). Compared to SM without hormone, ABA-treated *A. filiculoides* recovered comparably, *Azolla sp.* Anzali recovered without the cold-exposed red tissue withering, and *Azolla sp.* Bordeaux recovered better (**Figure 2B, Figure S2**). ABA could therefore boost other cold tolerance processes besides inhibiting DA-production.

A closer look at cold-treated *A. filiculoides* showed that DAs start accumulating at the shoot apex before the red hue over the sporophyte becomes apparent (**Figure S3**). Striking around the shoot apex were also the anthocyanic vacuolar inclusions (AVIs) in different sizes and colors: large ones in either black or red, outnumbered by small ones in purple (**Figure 2C-D, Figure S3**). AVIs warn that DA-biosynthesis/catabolism enzymes may be linked to membrane transport targeted to specific compartments inside the vacuole. In recovering plants, much more cells contained AVIs and lost their overall pigmentation (**Figure 2E, Figure S4**). The observations suggested that DA-accumulation differs per tissue and requires further analysis on higher resolution.

### *N. azollae* filaments disappear after prolonged cold-treatment, yet reappear in the newly outgrowing green fronds during recovery

*N. azollae* was followed in crushed shoot tips under the microscope. In the cold, the filament numbers decreased gradually during the first week, after which no filaments could be detected (**Figure S3**). The overall fluorescent signal faded after two weeks, and completely disappeared after three weeks (**Figure S3**). Upon return to 21°C, fluorescent filaments were observed after four days, coinciding with the first green outgrowth (**Figure S4**). After two weeks recovery at 21 °C, the overall filament numbers and fluorescent signal returned (**Figure S4**). The shoot apical *N. azollae* colony (SANC) must therefore survive the cold, possibly as single-cell akinetes.

After one week cold-treatment, leaf pocket (LP) isolation yielded some LPs full of red *N. azollae* filaments (**Figure 3A-B**), whilst others had no *N. azollae* filaments. We thus wondered where the DAs synthesized by the fern host accumulate.

**Figure 3.**
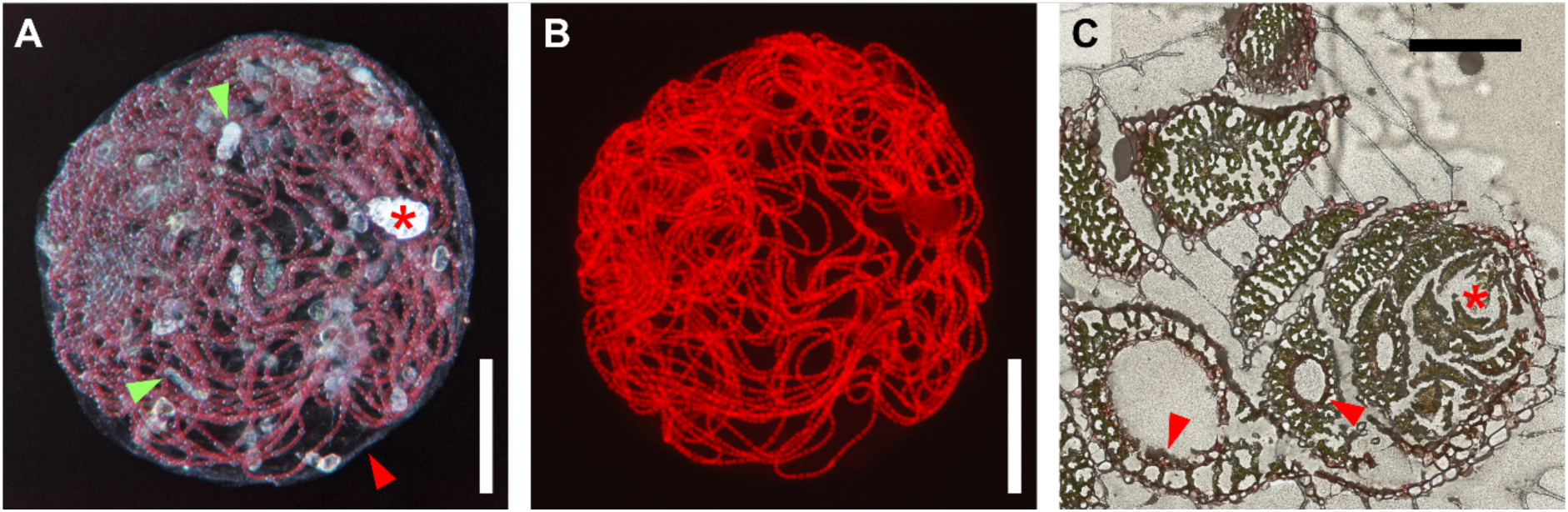
Red *N. azollae* and the tissue-specific localization of red pigmentation in Azolla. **A** Leaf pocket isolated from one-week cold-treated *A. filiculoides* under brightfield microscopy showing red *N. azollae* filaments, numerous translucent trichomes (green arrows), the surrounding membrane (red arrow) and the leaf pore (*). **B** Red filaments from **A** co-localize with the RFP autofluorescence of *N. azollae*. **C** Cross-section of three-weeks cold-treated *A. filiculoides* as used for LDI-MSI. Red pigment accumulation can be seen in the outer epidermis exposed to light, and in the epithelial cells surrounding the leaf cavities (arrows). The epidermis of the inner leaves around the shoot apex (*) and the parenchyma appear green. Scale bars: white and black, respectively, correspond to 100 µm and 200 µm.

### LDI-MSI reveals differential spatial localization of individual DA and CQA varieties in green and cold-treated red *A. filiculoides*

We embedded *A. filiculoides* after a three week cold-treatment in 20% gelatin, and prepared cryosections for further analysis. The parenchyma appeared green (**Figure 3C**). The red color mainly accumulated in the epidermis, and in the cells lining the LPs (**Figure 3C**). Surprisingly, the inner leaves with a green “outer” epidermis also had a red “inner” epidermis (**Figure 3C**). These latter cells originate from inward folding of the developing leaf and thus have an epidermal origin. Consistently, the LP trichomes and the outer papillae both stained for PAs (**Figure S5**).

To map the spatial distribution of individual molecules with LDI-MSI, we analogously prepared cryosections of red *A. filiculoides* after a three-week cold-treatment embedded together with untreated green sporophytes, for direct comparison. We sampled a region of interest (ROI) containing both red and green sporophyte tissue with a UV-laser at 20 µm pixel size, yielding a full mass spectrum per position. Any m/z feature detected could therefore be visualized in an image of the ROI, where each pixel represents an analyzed position.

The mass features *m/z* 293.0216 and 309.0165 could be putatively assigned to the [M-H+K]^+^ of the two DAs apigenidin and luteolinidin, respectively, and were verified with commercial standards (**Figure 4A-C**). High intensity pixels in the spatial distributions of these two compounds overlapped with the red regions in the optical image of the cryosection, and were not present in the green sporophyte (**Figure 4A-C**). Apigenidin was detected the most at the outermost tip of leaves, in the shoot apex and to a lesser extent throughout the parenchyma (**Figure 4B**). Luteolinidin was mostly detected in the outer epidermis and to a lesser extent in the cells lining the LPs (**Figure 4C**). Different types of Azolla DAs therefore have a different spatial distribution, and therefore, possibly, a different function. The mass feature *m/z* 615.4235 was repeatedly detected inside LPs of the red sporophyte, and was putatively assigned to the [M+K^+^]^+^ of 1-(O-α-D-glucopyranosyl)-(1,3R,25R)-hexacosanetriol (**Figure 4D**), a glycolipid previously extracted from *A. pinnata* (Ravi et al., 2020). This assignment is further corroborated by the presence of m/z 577.4678, which corresponds to the exact mass of the [M+H^+^]^+^ adduct ion of the proposed glycolipid. Similar glycolipids are reportedly incorporated in the cyanobacterial heterocyst envelope limiting entry of O_2_ (Soriente et al., 1992). Soriente et al. furthermore report the existence of congeners, where one of the hydroxyl groups is oxidized to a ketone. With the exception of the keto-analog, present as both [M+H^+^]^+^and [M+K^+^]^+^, m/z 575.4521 and 613.4081, respectively, we could not detect any other m/z features coinciding with the same spatial distribution.

**Figure 4.**
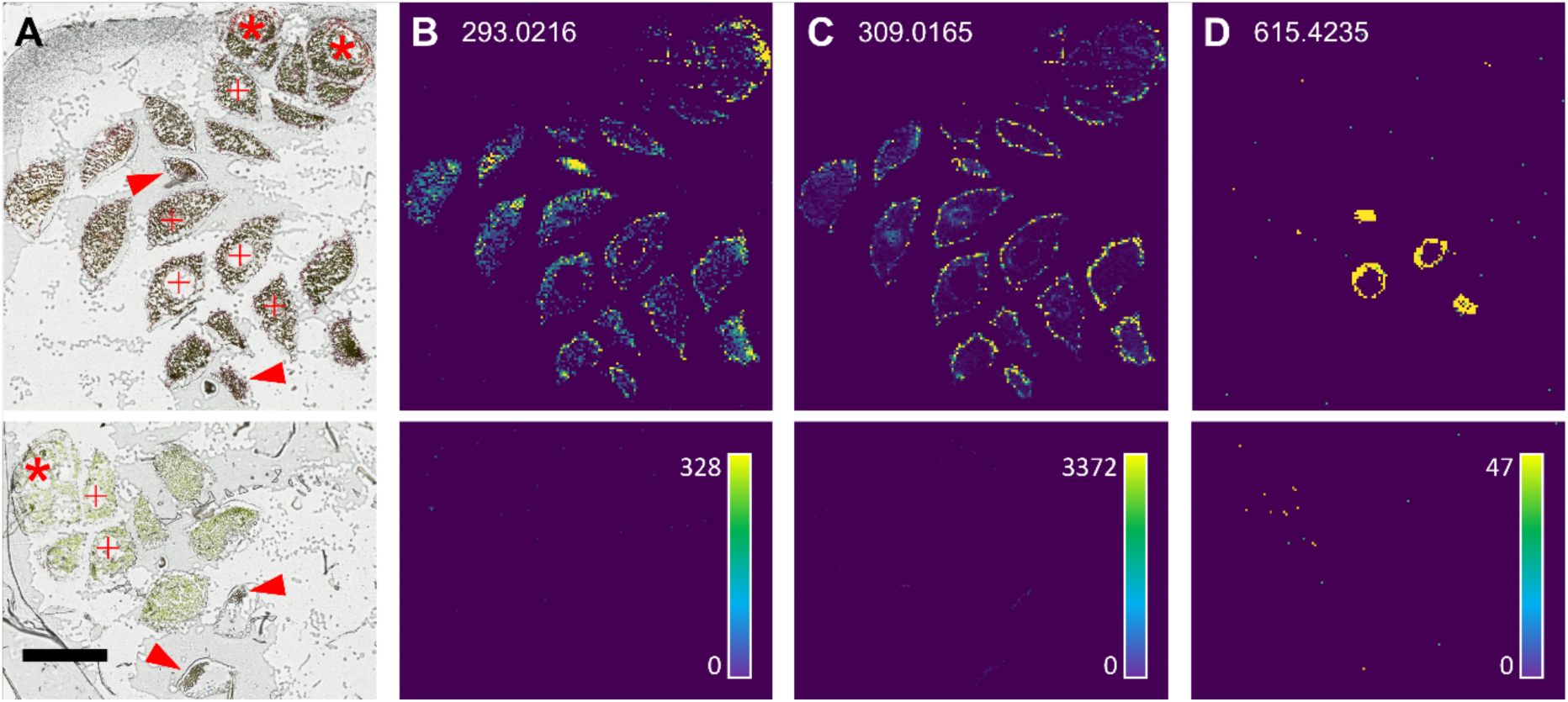
DA-related LDI-MSI maps of cold-treated (upper row) and untreated (lower row) *A. filiculoides*. Mass values were acquired from the same cryosection for direct comparison. Intensity scales are provided per mass value in the lower row. **A** The overview of the ROI. Indicated are shoot apices (*), leaf cavities (+) and leaf tips (arrows). The high-resolution image is available as **Figure S6**. Scale bar corresponds to 500 µm. Maps shown are for **B** *m/z* 293.0216 assigned to the [M-H+K]^+^ of apigenidin, **C** *m/z* 309.0165 assigned to the [M-H+K]^+^ of luteolinidin, and **D** *m/z* 615.4235 putatively assigned to the [M+K^+^]^+^ of 1-(O-α-D-glucopyranosyl)-(1,3R,25R)-hexacosanetriol.

We continued visualizing the spatial distribution of other key Azolla phenolics such as PAs and CQAs. *Azolla* PAs have an average degree of polymerization of seven and exceed the molecular weight limit of detection. Instead, we chose to focus on three different PA mono– and dimers present in Azolla, as indicators (Güngör et al., 2021). The mass features *m/z* 329.0416, 615.0887, and 617.1045 were putatively assigned to the [M+K^+^]^+^ of (epi)catechin, procyanidin A, and procyanidin B, respectively, and verified with commercial standards (**Figure 5A-D**). All three molecules were detected throughout the parenchyma and most intensely at the shoot apex of the red sporophyte, but they were more widespread and not restricted to the red parts, such as apigenidin (**Figure 4B**, **Figure 5A-D**). In the green sporophyte, PAs were only detected in the outermost tip of some leaves (**Figure 5A-D**). The mass feature *m/z* 363.0681 was putatively assigned to the [M+Na^+^]^+^ of aesculin, and was verified with a commercial standard (**Figure 5E**). This coumarin was also detected in leaf tips of the green sporophyte and throughout the parenchyma of the red sporophyte, but in contrast to PAs, it was more prevalent in older leaves instead of the shoot apex (**Figure 5A-E**). The mass features *m/z* 393.0574 and 555.0888, were putatively assigned to the [M+K^+^]^+^ of CQA and a diCQA, respectively (**Figure 5F-G**). Comparable to DAs, CQAs were detected throughout the parenchyma of the red sporophyte, exhibiting the highest signal intensity at the shoot apex, whilst diCQAs appeared to be highly localized in the outer epidermis (**Figure 5F-G**).

**Figure 5.**
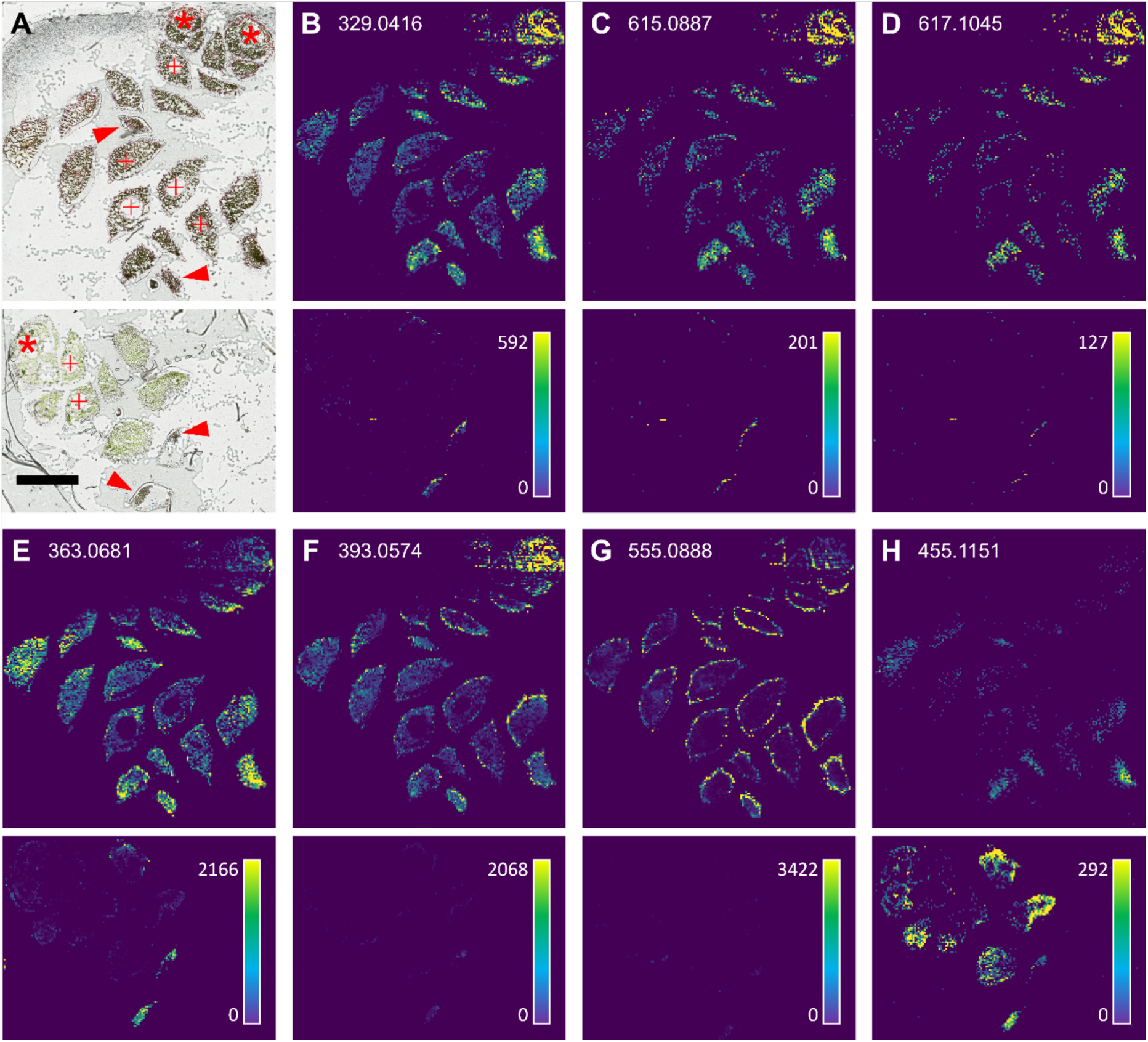
LDI-MSI maps of abundant Azolla phenolics in cold-treated (upper row) and untreated (lower row) *A. filiculoides*. M/z values were acquired from the same cryosection for direct comparison. Intensity scales are provided per mass value in the lower row. **A** Overview of the ROI as per Figure 4A. Scale bar corresponds to 500 µm. Maps shown are for **B** *m/z* 329.0416 assigned to the [M+K^+^]^+^ of (epi)catechin, **C** *m/z* 615.0887 assigned to the [M+K^+^]^+^ of procyanidin A, **D** *m/z* 617.1045 assigned to the [M+K^+^]^+^ of procyanidin B, **E** *m/z* 363.0681 assigned to the [M+Na^+^]^+^ of aesculin, **F** *m/z* 393.0574 putatively assigned to the [M+K^+^]^+^ of caffeoylquinic acid, **G** *m/z* 555.0888 putatively assigned to the [M+K^+^]^+^ of dicaffeoylquinic acid, and **H** *m/z* 455.1151 putatively assigned to the [M+Na^+^]^+^ of hypothetic compound C_18_H_24_O_12_.

Generally, m/z features present in both phenotypes exhibited higher intensities in the cold-treated red sporophyte (**Figure 4 and 5**). However, some m/z features exhibited the opposite trend, for example, *m/z* 455.1151, putatively assigned to the [M+Na^+^]^+^ of C_18_H_24_O_12_, based on accurate mass and isotopic pattern, but no reasonable structure was putatively assignable with the information at hand (**Figure 5H**). While homogeneously distributed at higher signal intensities throughout the tissue of the green sporophyte, almost no signal intensity was observed in the areas correlating with the tissue of the red sporophyte (**Figure 5H**).

### The *N. azollae* transcript profile during cold acclimation differs from that of akinetes in sporocarps

Transcript accumulation in host and endosymbiont were acquired simultaneously (dual RNA-seq) to profile the symbiosis physiological response. *A. filiculoides* sporophytes were grown at 21°C and 4°C to yield the samples untreated day 14 (**U14**), cold-treated day 14 (**C14**) and cold-treated day 21 (**C21**) (**Figure S7**). Reads mapping on the *N. azollae* genome were drastically reduced after cold-treatment (**Figure S8**), consistent with absence of *N. azollae* filaments after prolonged cold-treatment (**Figure S3**); yet the sequencing depth sufficed for differential gene expression analysis with DESeq2.

822 significantly (padj<0.1) differentially expressed genes overlapped between **C14** and **C21**, respectively compared to **U14** (**File S1**). The gene amongst those with the highest transcript accumulation was histidinol dehydrogenase (AAZO_RS04970) which together with the high HisF (AAZO_RS09015) transcript indicated up-regulated histidine production (**Table 1**). Abundant transcripts of form 1 RuBisCo large subunit (AAZO_RS10205) and phosphoribulokinase (AAZO_RS23080) were downregulated, along with the Nif-operon, making photosynthesis, CO_2_-fixation and N_2_-fixation decreased processes in cold-treated *N. azollae* in terms of transcript investment (**Table 1**).

**Table 1.** *N. azollae* genes of key processes with differentially accumulating transcripts (padj<0.1) during cold-treatment. **C14** and **C21** were, respectively, compared to **U14** (untreated, **U14**; 14-days cold-treated, **C14**; and 21-days cold-treated, **U21**). Source data in **File S1**.

| Gene | Function (NCBI) | C14 versus U14 |  | C21 versus U14 |  |
| --- | --- | --- | --- | --- | --- |
|  |  | baseMean | log2Fold Change | baseMean | log2Fold Change |
| Histidine production |  |  |  |  |  |
| AAZO_RS04970 | histidinol dehydrogenase | 17289 | 0.61 | 18316 | 0.96 |
| AAZO_RS09015 | HisF | 394 | 2.46 | 214 | 1.7 |
| Photosynthesis and CO<sub>2</sub>-fixation |  |  |  |  |  |
| AAZO_RS10205 | form I RuBisCo large subunit | 12680 | -2.3 | 13704 | -2.44 |
| AAZO_RS23080 | phosphoribulokinase | 12163 | -2.27 | 14413 | -1.49 |
| Nif-operon |  |  |  |  |  |
| AAZO_RS06465 | NifW | 67 | -6.17 | 78 | -4.38 |
| AAZO_RS06480 | NifX | 558 | -5.96 | 647 | -5.03 |
| AAZO_RS06485 | NifN | 1252 | -3.41 | 1356 | -4.76 |
| AAZO_RS06490 | NifE | 478 | -3.54 | 573 | -2.24 |
| AAZO_RS06495 | NifK | 1211 | -2.75 | 1321 | -4.21 |
| AAZO_RS06500 | NifD | 4046 | -2.19 | 4264 | -2.98 |
| AAZO_RS06505 | NifH | 1758 | -3.63 | 1971 | -3.91 |
| AAZO_RS06510 | NifU | 1530 | -3.88 | 1787 | -3.29 |
| AAZO_RS06515 | NifS | 3748 | -3.1 | 4264 | -2.73 |
| AAZO_RS06525 | NifB | 2213 | -2.52 | 2430 | -2.86 |
| AAZO_RS06575 | NifZ | 15 | -4.24 | 17 | -2.71 |
| AAZO_RS06580 | NifV | 707 | -1.32 | 658 | -2.79 |
| Photooxidative damage mitigation |  |  |  |  |  |
| AAZO_RS34525 | ssl1498 family light-harvesting-like protein | 618 | 4.84 | 277 | 3.88 |
| AAZO_RS13445 | chlorophyll a/b-binding protein | 459 | 4.2 | 353 | 4.03 |
| AAZO_RS12085 | orange carotenoid protein | 1335 | 0.86 | 1669 | 1.56 |
| AAZO_RS19455 | HAMP domain-containing sensor histidine kinase | 982 | 3.63 | 586 | 3.22 |
| AAZO_RS03300 | ATP-dependent Clp protease adapter ClpS | 218 | 1.19 | 176 | 0.96 |
| AAZO_RS04880 | ATP-dependent Clp protease ATP-binding subunit | 10859 | 2.65 | 9863 | 2.81 |
| AAZO_RS14105 | ATP-dependent Clp protease proteolytic subunit | 324 | 2.01 | 284 | 1.97 |
| AAZO_RS13760 | chaperonin GroEL | 3282 | 2.45 | 1901 | 1.73 |
| AAZO_RS09690 | molecular chaperone DnaK | 831 | 3.59 | 818 | 3.87 |
| AAZO_RS15995 | molecular chaperone DnaK | 1036 | 1.44 | 1188 | 1.95 |
| AAZO_RS21990 | Peroxiredoxin | 11300 | 2.23 | 7516 | 1.77 |
| AAZO_RS21995 | redoxin domain-containing protein | 704 | 2.13 | 680 | 2.42 |
| AAZO_RS22000 | glutathione S-transferase family protein | 149 | 2.34 | 188 | 3.17 |
| Cell division and differentiation |  |  |  |  |  |
| AAZO_RS05815 | FtsE | 56 | -3.08 | 60 | -4.07 |
| AAZO_RS09060 | FtsZ | 921 | -1.72 | 1016 | -1.47 |
| AAZO_RS17725 | HetR | 411 | 1.2 | 400 | 1.45 |

The most upregulated genes were ssl1498 family light-harvesting-like protein (AAZO_RS34525) and chlorophyll a/b-binding protein (AAZO_RS13445) (**Table 1**), both high-light-inducible proteins and therefore, most likely, mitigating photooxidative damage caused by cold (Dittami et al., 2010; Rahimzadeh-Karvansara et al., 2022). An orange carotenoid protein (OCP, AAZO_RS12085) was also upregulated (**Table 1**). OCP binds to phycobilisomes to quench excess energy (Thurotte et al., 2015); it is orange upon production but turns red upon photoactivation, which could possibly explain the red *N. azollae* filaments observed in leaf pockets from cold-acclimating symbioses (**Figure 3A-B**).

Response to photooxidative stress was further supported by upregulation of a sensor histidine kinase (AAZO_RS19455) (**Table 1**), belonging to the same orthogroup as Hik34 from *Synechocystis sp.* PCC 6714 (OrthoDB v11, orthogroup 122199at1117, accessible via www.orthodb.org). The Hik34-ortholog furthermore may have caused up-regulation of the transcripts encoding the ATP-dependent Clp protease (AAZO_RS03300, AAZO_RS04880 and AAZO_RS14105), heat-shock-protein GroEL (AAZO_RS13760) and DnaK (AAZO_RS09690 and AAZO_RS15995) (**Table 1**); the latter transcripts are known to be involved in cyanobacterial cold tolerance (Porankiewicz et al., 1998; Hossain & Nakamoto, 2002). Upregulation of a cluster of peroxiredoxin (AAZO_RS21990), redoxin domain-containing protein (AAZO_RS21995) and glutathione S-transferase family protein (AAZO_RS22000) furthermore pointed towards an increased maintenance of the cell redox state (**Table 1**).

Downregulation of cell division protein FtsE and FtsZ (AAZO_RS05815, AAZO_RS09060) (**Table 1**), made us wonder whether *N. azollae* survives the cold as single-cell akinetes at the shoot apex. Comparison of the 822 cold-related genes with the 1089 significantly (padj<0.1) differentially expressed *N. azollae* genes in megasporocarps (**mega** compared to **sp**), resulted in 237 overlapping genes (**File S1**). The minimal overlap between the differential expression patterns of those genes between the cold samples and the megasporocarp, however, indicated that the *N. azollae* physiology after prolonged cold is very different to that of akinetes from the megasporocarp (**Figure S9**). The heterocyst differentiation master regulator HetR (AAZO_RS17725) was for example upregulated in cold, while downregulated in the megasporocarp (**Table 1, File S1**).

### Fern phenolic biosynthesis enzymes and potential regulators during prolonged cold-treatment of *A. filiculoides*

Relative amounts of transcripts the fern host invested per metabolic process, as defined by automatic Mercator annotation of Afi_v2 genes (Lohse et al., 2014; Güngör et al., 2023), after 14 days (**C14**) and 21 days (**C21**) cold-treatment were compared to the untreated control after 14 days (**U14**). Photosynthesis (239 genes) was the most reduced process, while secondary metabolism (60 manually curated genes) was the most induced (**Figure S10; File S2**), consistent with literature (Tran et al., 2020). 1943 significantly (padj<0.1) differentially expressed genes overlapped between **C14** and **C21**, respectively compared to **U14** (**File S2**). The gene amongst those with the highest transcript accumulation was an early light-induced protein (ELIP, Afi_v2_s37G001720.1) (**Table 2**); likely mitigating photooxidative damage, consistent with what is known of plant acclimation to cold (Heddad et al., 2006). An upregulated allene oxidase synthase (AOS, Afi_v2_s16G003320.1) and 12-oxophytodienoate reductase (OPR, Afi_v2_s31G000290.1) indicated increased production of jasmonic acid (**Table 2**). Two ammonium transporters (AMT, Afi_v2_s40G002100.1 and Afi_v2_s41G000920.1) were also upregulated (**Table 2**), indicative of an altered source of ammonium in line with reduced N_2_-fixation by the cyanobacterial endosymbiont.

**Table 2.** *A. filiculoides* genes with differential transcript accumulation (padj<0.1) during cold-treatment and from categories discussed in the text. **C14** and **C21** were, respectively, compared to **U14** (untreated, **U14**; 14-days cold-treated, **C14**; and 21-days cold-treated, **U21**). Source data in **File S2**.

|  |  | C14 versus U14 |  | C21 versus U14 |  |
| --- | --- | --- | --- | --- | --- |
| Gene | Function (Mercator) | baseMean | log2Fold Change | baseMean | log2Fold Change |
| Photooxidative damage mitigation |  |  |  |  |  |
| Afi_v2_s37G001720.1 | ELIP LHC-related protein | 42581 | 3.47 | 42315 | 3.51 |
| Jasmonic acid production |  |  |  |  |  |
| Afi_v2_s16G003320.1 | allene oxidase synthase (AOS) | 449 | 1.25 | 435 | 1.25 |
| Afi_v2_s31G000290.1 | 12-oxophytodienoate reductase (OPR) | 8169 | 2.76 | 4372 | 1.74 |
| Ammonium transporters |  |  |  |  |  |
| Afi_v2_s40G002100.1 | ammonium transporter (AMT2/3) | 284 | 2.63 | 163 | 1.67 |
| Afi_v2_s41G000920.1 | ammonium transporter (AMT2/3) | 497 | 2.46 | 216 | 0.91 |
| Transcription factors |  |  |  |  |  |
| Afi_v2_s1G000550.2 | MYB (FLP) | 732 | 1.67 | 613 | 1.39 |
| Afi_v2_s29G001620.1 | bHLH | 572 | 1.46 | 563 | 1.57 |
| Afi_v2_s2495G000440.1 | MYB (VIII-D) | 539 | 0.65 | 398 | 0.13 |
| Afi_v2_s25G003460.1 | MYB (VIII-D) | 707 | 0.17 | 1065 | 1.14 |
| Afi_v2_s42G000270.4 | MYB (VIII-D) | 557 | 0.20 | 569 | 0.34 |
| PBP/FBP-enzymes |  |  |  |  |  |
| Afi_v2_s37G002190.1 | Phenylalanine ammonia lyase (PAL) | 6976 | 0.54 | 6532 | 0.48 |
| Afi_v2_s15G004580.2 | Phenylalanine ammonia lyase (PAL) | 783 | 1.16 | 925 | 1.63 |
| Afi_v2_s3G002840.2 | Phenylalanine ammonia lyase (PAL) | 2516 | 2.15 | 1687 | 1.46 |
| Afi_v2_s1G000760.3 | Phenylalanine ammonia lyase (PAL) | 6870 | 1.00 | 5867 | 0.74 |
| Afi_v2_s12G003500.2 | Cinnamate 4-monooxygenase (C4H) | 28394 | 2.53 | 27519 | 2.53 |
| Afi_v2_s54G001950.8 | 4-Coumarate-CoA ligase (4CL) | 1825 | 0.49 | 1503 | 0.11 |
| Afi_v2_s37G000480.1 | Chalcone synthase (CHS) | 29707 | 0.91 | 29831 | 1.00 |
| Afi_v2_s41G002720.1 | Chalcone synthase (CHS) | 8560 | 1.71 | 5825 | 1.00 |
| Afi_v2_s72G000900.1 | Chalcone synthase (CHS) | 14444 | 2.02 | 15383 | 2.19 |
| Afi_v2_s26G004110.1 | Chalcone isomerase (CHI) | 1183 | 0.57 | 1068 | 0.45 |
| Afi_v2_s74G000210.2 | Leucoanthocyanidin reductase (LAR) | 3073 | 1.11 | 3610 | 1.52 |
| Most upregulated |  |  |  |  |  |
| Afi_v2_s29G003930.6 | GDSL esterase/lipase | 17580 | 8.16 | 10858 | 7.49 |
| Afi_v2_s35G001880.1 | Cytochrome P450 CYP736A12 | 220 | 8.02 | 250 | 8.20 |
| Afi_v2_s12G002030.1 | AfDFR1 | 4317 | 8.14 | 3022 | 7.67 |

A single MYB TF (Afi_v2_s1G000550.2) and five bHLH TF were cold-induced; one bHLH TF (Afi_v2_s29G001620.1) had transcript levels comparable to the MYB (**Table 2**). Based on phylogenetic analyses by Jiang & Rao, 2020, this MYB is FLP-type (**File S2**). Stress-induced flavonoid-accumulation in angiosperms was attributed to VIII-E-type MYBs, which had negligible transcript accumulation in cold-treated *A. filiculoides* (**File S2**).

Homologs of the three core enzymes of the PBP (PAL, C4H and 4CL) and the first two enzymes of the FBP (CHS and CHI) were all upregulated in **C14** but in **C21** some decreased in expression, whilst others continued increasing (**Table 2**). Leucoanthocyanidin reductase (LAR, Afi_v2_s74G000210.2), involved in PA-biosynthesis (Güngör et al., 2021), also continued increasing (**Table 2**). Candidate enzymes participating in CQA-production (HCT/HQT, C3H and CSE), identified with Mercator and verified based on phylogenetic analyses by (de Vries *et al*. 2021), had varying differential expression (**File S2**).

Amongst the top 10 most upregulated genes in cold, that could potentially be involved in phenolic metabolism, were a GDSL esterase/lipase protein (GELP, Afi_v2_s29G003930.6), a cytochrome-P450 enzyme (CYP450, Afi_v2_s35G001880.1) and a bifunctional dihydroflavonol 4-reductase/flavanone 4-reductase (DFR1, Afi_v2_s12G002030.1) (**Table 2**). The latter had negligible transcript accumulation in green ferns (**U14**), but reached transcripts levels of PBP/FBP-enzymes in cold-treated red ferns (**Table 2**). As candidate first enzyme for DA-biosynthesis its regulated expression could thus act as a gatekeeper (**Figure 1B**). We therefore studied *Af*DFR1 in more detail.

### *Af*DFR1 enzyme converts the flavanone substrates naringenin and eriodictyol *in vitro*

We identified 13 candidates with BLAST-searches through Afi_v2 genes with sequences from the *Sb*FNR orthogroup (Sobic.006G226800), extracted from the 1KP database (Wong et al., 2020). Mercator automatically annotated five as DFR, five as tetraketide α-pyrone reductase (TKPR) and three as cinnamoyl-CoA reductase (Lohse et al., 2014). Based on phylogenetic analysis, only four of the five DFRs seemed related (**Figure S11**). Only *Af*DFR1 (Afi_v2_s12G002030.1) and a TKPR (Afi_v2_s5G008050.13) were upregulated in cold, but the latter was already expressed in green ferns and therefore unlikely related to DA-accumulation (**Figure 6A**).

**Figure 6.**
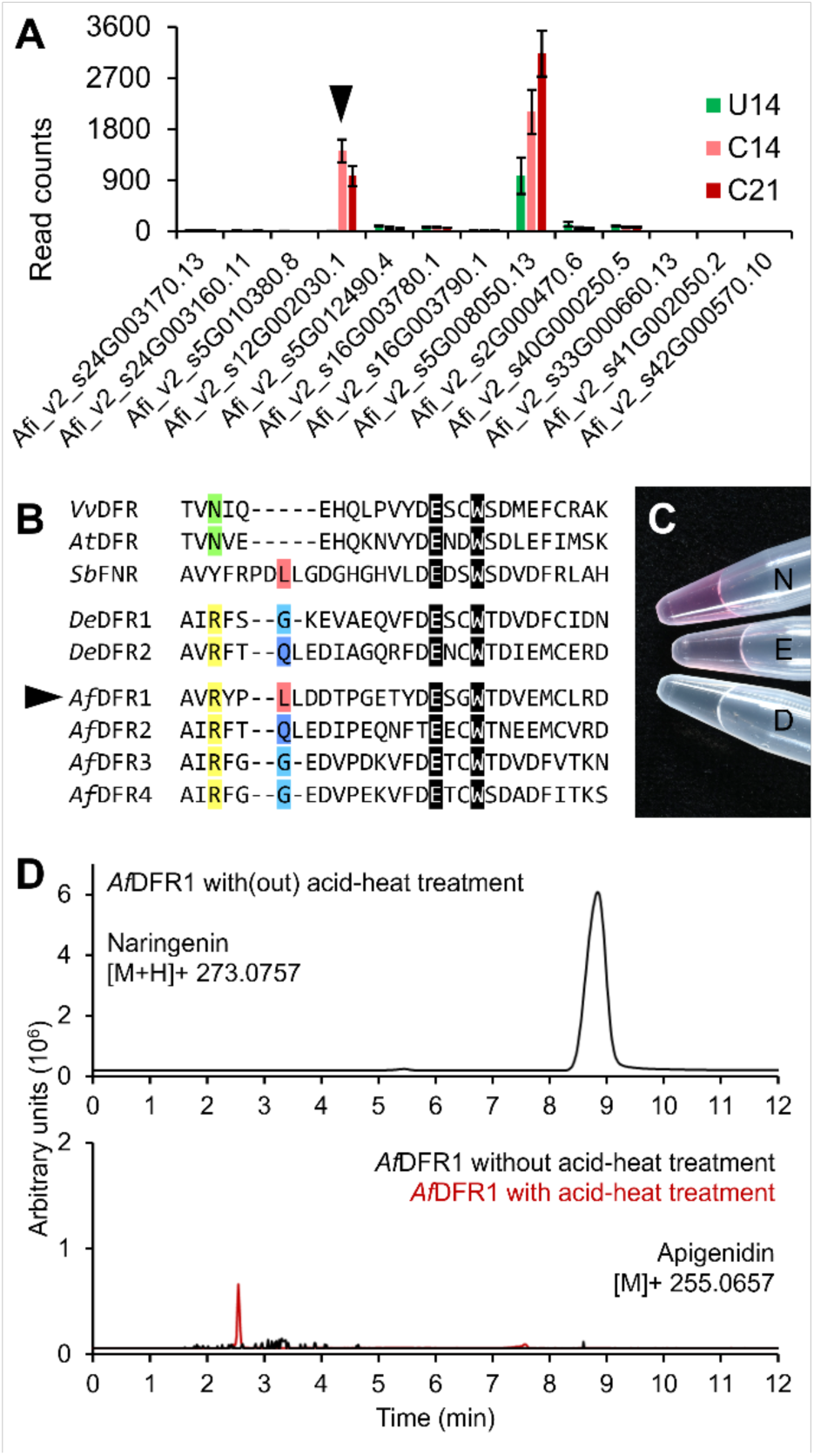
*Af*DFR1 expression pattern, substrate-interaction region and invitro enzyme assay. **A** Transcript accumulation of 13 DFR-related genes found in the Afi_v2 genome in untreated (**U14**), 14-days cold-treated (**C14**) and 21-days cold-treated (**U21**) *A. filiculoides*. Arrow indicates *Af*DFR1. Standard deviations are for n=3. **B** Alignment of the 26-amino-acid-long substrate-interaction region of *V. vinifera* DFR (Petit et al., 2007), with that of *A. thaliana* DFR, *S. bicolor* FNR, two *D. erythrosa* DFRs (Chen et al., 2020) and the four *A. filiculoides* DFRs (**Figure S11**). Color-marked are shared AAs of interest. **C** Color-development after acid-heat treatment of the *Af*DFR1 enzyme assay reaction with naringenin (N), eriodictyol (E) and dihydroquercetin (D) as a substrate. **D** LC-MS analysis of the naringenin assay from **C**, the assay was split in two after the reaction so as to have one without and one with acid-heat treatment. Top: the naringenin peak was identical. Bottom: the apigenidin peak appeared only after acid-heat treatment consistent with color-development.

To identify possible motifs for substrate specificity, the region encompassing amino-acids interacting with the substrate in the *Vitis vinifera* DFR was aligned with that of *At*DFR, *Sb*FNR, two *Dryopteris erythrosa* DFRs and the four *Af*DFRs (Petit et al., 2007; Chen et al., 2020). *Sb*FNR and the fern sequences had 3-5 amino-acid-long insertions in the active site which we named L-type, G-type and Q-type (**Figure 6B**). *De*DFR2 could substitute *At*DFR in the *A. thaliana tt3* DFR-knockout-mutant whilst *De*DFR1 could not (Chen et al., 2020). Q-type DFRs, therefore, likely participate in PA-production. L-type *Sb*FNR and *Af*DFR1 shared the amino acids proline, leucine and aspartic acid at the insertion site (**Figure 6B**). *Af*DFR1 could therefore likely convert flavanones into flavan-4-ols (**Figure 1B**).

To assay *Af*DFR1, we fused its gene sequence to a N-terminal polyhystidine-tag and included 35S::GFP and kanamycin resistance as reporter genes in the plasmid. We could detect GFP-signal 72 hours after Agro-infiltrating *N. tabacum* leaves (**Figure S12**). Assuming *Af*DFR1 would be expressed at the same time as GFP, as both were driven by the same promotor, we choose this timepoint for our assay. We used the substrates naringenin (N), eriodictyol (E) and dihydroquercetin (D). When the enzyme was purified on a Ni-NTA column, only naringenin and eriodictyol resulted in color formation after acid-heat treatment (**Figure 6C**). The same procedures with a mock-plasmid with only the reporter genes never resulted in color formation. We therefore attributed the color formation to the activity of *Af*DFR1 and concluded it can only use flavanones *in vitro* consistent with FNR-activity. As naringenin gave the most intense color indicative for the best conversion rate, we ran this sample on the LC-MS for product confirmation. Without acid-heat treatment only naringenin was detected whilst after acid-heat treatment also apigenidin could be detected (**Figure 6D**). The expected *m/z* for the flavan-4-ol intermediate apiforol was not detected.

To test whether *Af*DFR1 can substitute *At*DFR *in planta*, we transformed *Af*DFR1 in *A. thaliana tt3* mutants. Three transgenics were selected on antibiotics and GFP-expression in the leaves was verified. The plants did not accumulate red pigments under high light intensity as opposed to WT Col-0 *A. thaliana*, and the harvested seeds remained yellow (data not shown). *Af*DFR1 can therefore not substitute *At*DFR and restore HA-production in *A. thaliana*, consistent with the results of the *in vitro* enzyme assay (**Figure 6C**). *Af*DFR1 thus likely participates in DA-biosynthesis and defines the substrate flux into this pathway whilst *A. thaliana* does not possess, or at least express, the required additional factors for DA-production (**Figure 1B**).

The second and final enzymatic step from flavan-4-ols to 3-deoxyanthocyanidins is generally attributed to ANS (**Figure 1B**). ANS belongs to the 2-oxoglutarate dependent dioxygenase (2OGD) superfamily, for which we calculated a phylogenetic tree including 29 identified *A. filiculoides* 2OGDs using the 1KP orthogroup database in Güngör et al., 2023. Clades with flavonoid-related 2OGDs were combined into a new pruned tree and included 7 *A. filiculoides* 2OGDs (**Figure 7A**). The tree showed that flavonol synthase (FLS) and ANS diverged in seed plants whilst *A. filiculoides* has an FLS/ANS-like gene (Afi_v2_s19G002650.1). *Af*FLS/ANS-like expression was downregulated in cold and this enzyme is thus unlikely required for DA-production (**Figure 7B**).

**Figure 7.**
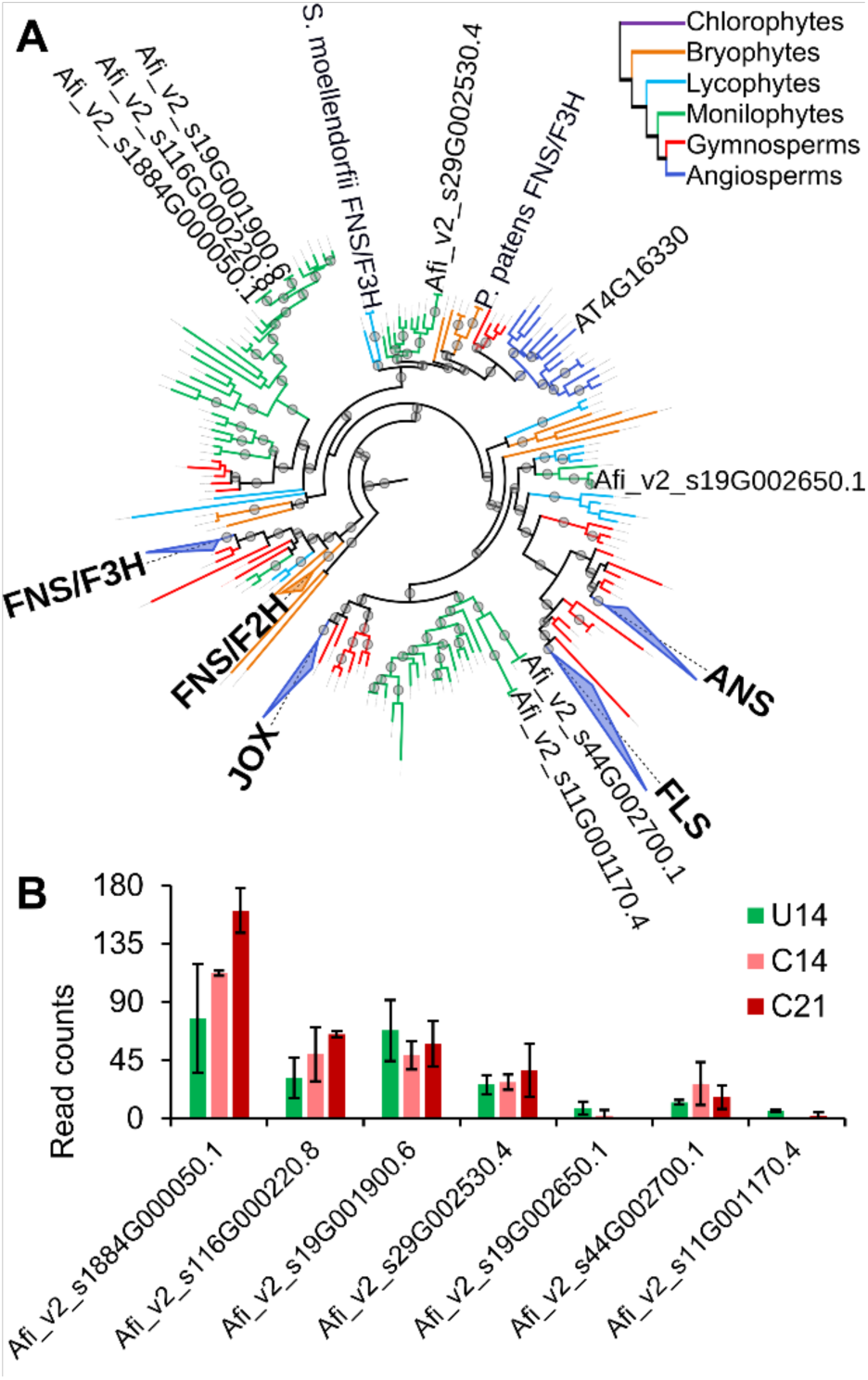
Phylogeny and expression pattern of Azolla 2OGD-enzymes related to the FBP. **A** Pruned version of the 2OGD-superfamily orthogroup-based phylogenetic tree from Güngör et al., 2023, showing enzymes related to the FBP, including 7 *Af*2OGDs and functionally characterized 2OGDs from angiosperms, lycophytes and bryophytes (Li et al., 2020). Branches with grey circles have more than 80% bootstrap support. Enzymes, FNS: flavone synthase, F2H: naringenin 2-hydroxylase, F3H: flavanone 3-hydroxylase, ANS: anthocyanidin synthase, FLS: flavonol synthase, and JOX: jasmonic acid oxidase. **B** Transcript accumulation of the 7 *Af*2OGDS from **A** in untreated (**U14**), 14-days cold-treated (**C14**) and 21-days cold-treated (**U21**) *A. filiculoides*. Standard deviations are for n=3.

## Discussion

### High resolution spatial analysis suggests distinct functions for individual members of the same phenolic class

Phenolics can have different activities and functions depending on hydroxylation, glycosylation, methylation and acylation (Alseekh et al., 2020). 5-Methoxyluteolinidin, for example, exhibits more antifungal properties than luteolinidin (Lo et al., 1996). We therefore compared the spatial distribution of the abundant apigenidin and luteolinidin, which differ in the hydroxylation of position 3’ of the B-ring (**Figure 1B**). Apigenidin was more predominant in the leaf tips and parenchyma whilst luteolinidin was more epidermis-specific (**Figure 4A-C**). Apigenidin could therefore be scavenging ROS in the photosynthetically active parenchyma, whilst luteolinidin may prevent blue/green light from reaching the parenchyma and, in addition, inhibit microbial infection of the plant surface by specifically inhibiting tyrosinase activity (Panis et al., 2021; Yang et al., 2022).

PAs are most known as antifeedant and the mono-/dimeric subunits exhibited the highest intensities throughout the parenchyma, with maxima detected in the youngest leaves at the shoot apex (**Figure 5A-D**). PAs might therefore possibly increase in degree of polymerization as the leaves age. Similarly, aesculin, known for its bitter taste (Ribéreau-Gayon et al., 2021), was detected in the parenchyma but exhibited maximal intensities in adult leaves (**Figure 5E**). CQAs, on the other hand, are most known for their ROS-scavenging and antimicrobial capacities (Mondolot et al., 2006). CQAs exhibited a similar spatial distribution to PAs, but diCQAs were highly localized in the outer epidermis (**Figure 5F-G**). It therefore seems that PAs and aesculin, phenolics deterring big grazers swallowing sporophytes as a whole, accumulate throughout the parenchyma, whilst those that protect from microbial infections are epidermis-specific (diCQAs).

LDI-MSI is especially suitable for analyzing spatial distributions of melecules in small plants like Azolla because all organs fit in one cryosection, promoting comparability between different tissue and shortening acquisition time, while the aromatic nature of the flavonoids make them well suitable for direct LDI analysis, as opposed to matrix-assisted LDI required for non-UV active compounds. The investigation of spatial distributions of individual molecules helped form hypotheses on their function and will improve our understanding of plant secondary metabolism. One limiting factor in the current state of the art methodologies is sensitivity (Knochenmuss, 2013; Potthoff et al., 2020). Generally, signal intensities observed in the cold-treated red sporophyte were higher than the signal observed for the untreated green sporophyte (**Figure 4 and 5**). This was evident for DAs induced by cold, but phenolics such as PAs and CQAs have always been detected in high concentrations in green Azolla sporophyte extracts (Qian et al., 2020; Güngör et al., 2021). Cold-treated plants might therefore accumulate even more PAs and CQAs, in line with most of the gene expression (**Table 2**), and dominate the signal. A specific study, linking signal intensities acquired in LDI-MSI experiments with absolute quantification of specific phenolic compounds from the very same plant individual, would be required to elucidate the causality. Any difference in the observed signal intensities between the two phenotypes may be compounded further by the fact that phenolics with aromatic rings serve as endogenous matrix. An increased concentration of phenolics might therefore provide more charge transfer capabilities, leading to higher ionization efficiencies (Wyplosz, 2003; Hölscher et al., 2009; Li et al., 2012).

### Azolla symbiosis partners during cold acclimation experience light-induced oxidative stress and are likely metabolically uncoupled

Gene expression profiles of both phototrophic organisms in the symbiosis reflected high levels of oxidative stress, likely due to the absorption of photons in the blue/green wavelength. DA-accumulation in light-exposed tissue of the fern host further supported this; DAs are known to absorb ROS as well as filter blue/green light and scatter red light. *N. azollae*, therefore, received excess red light when the fern host accumulated DA, which is typically harvested by its phycocyanobilin giving *N. azollae* their typical blue/green color. The red *N. azollae* in leaf pocket preparations of cold-acclimating symbioses (**Figure 3A**) was not due to phycoerythrobilins because they are encoded in the *N. azollae* genome (they harvest blue/green light; Six et al., 2007). Instead, reduced cell numbers and reads for photosynthesis, and high OCP protein transcripts (**Table 1; File S1**) were consistent with diminishing red-fluorescence and biomass, and red color upon leaf pocket preparation of *N. azollae* in the cold.

Transfer of a substantial amount of luteolinidin from AVIs to the LP through the cutinic cuticle of the LP seemed unlikely (Genet et al., 2002), but cannot be ruled out via exocytosis. Also, the LP trichomes, which are most likely essential for the symbiotic interaction (Zheng et al., 2008), were colorless (**Figure 3A**). Nevertheless, luteolinidin from the host may be an important intermediate in the biosynthesis of signaling compounds during physiological adaptation of the symbiosis to cold: Azolla DAs, including acetylated/glycosylated luteolinidin reduced expression of hormogonia factors in *N. punctiforme*, related to *N. azollae* (Cohen et al. 2002a). Almost all pilus/twitching genes of *N. azollae* associated with the formation of motile filaments were indeed downregulated in cold-treated symbioses (**File S1**).

A glycolipid generally found in the cyanobacterial heterocyst envelope (Soriente et al., 1992), was detected in LPs of cold-treated *Azolla* (**Figure 4D**). The *N. azollae* glycolipid gene cluster (Hgl-island) may be substantially translated after cold-treatment. HglB encodes a polyketide synthase required for the production of both heterocyst and akinete glycolipids; and akinetes of mutants lacking HglB were more vulnerable to freezing (−20°C) in free-living *Nostoc* sp. (Garg & Maldener, 2021b).

Glycolipids could therefore be essential for *N. azollae* cold acclimation, and their production could possibly be driven by HetR (Wolk et al., 1994; Khudyakov & Golden, 2004). This was consistent with the transcript accumulation of HetR that specifies heterocyst differentiation, in spite of the low accumulation of Nif-operon transcripts, in *N. azollae* (**Table 1**), and upregulation of AMTs in the fern host (**Table 2**) of cold-treated symbioses. HetR upregulation could also be linked to the extreme accumulation of histidinol dehydrogenase (**Table 1**), required for the assembly of heterocyst cell walls (Lechno-Yossef et al., 2011). Without its own N_2_-fixation, the endosymbiont may depend on the host for its nitrogen supply. High transcript accumulation of histidinol dehydrogenase may thus reflect decoupling of metabolism from host and *N. azollae* during symbiosis cold acclimation, as in the case of the legume-rhizobia symbiosis (Yadav et al., 1998).

### The first DA-biosynthesis enzyme *Af*DFR1 is only transcribed when the symbiosis is exposed to cold

Expression and *in vitro* enzyme assays indicate that *Af*DFR1 is the first enzyme of Azolla DA-biosynthesis (**Figure 6**). As in Sorghum, *Af*DFR1 transcripts only accumulated in the cold, suggesting that the DA pathway is regulated transcriptionally in ferns and in those seed plants that can produce DAs. Surprisingly, VIII-E-type MYBs, responsible for stress-induced flavonoid-accumulation in seed plants (Jiang & Rao, 2020), were not expressed in cold-treated Azolla indicating that the regulation of DAs is carried out by different TFs and integrated differently in the ferns. Consistently, in *A. thaliana* ABA and JA induced HA-biosynthesis (Loreti et al., 2008), whereas in Azolla ABA inhibited DA-accumulation (**Figure 2A**). The most-upregulated MYB in cold was FLP-type (Afi_v2_s1G000550.2) (**Table 2**), known for the regulation of stomatal development in *A. thaliana* (Xie et al., 2010). Overexpression of 35S::Afi_v2_s1G000550.2 in *A. thaliana* and *N. tabacum* did not yield an apparent phenotype in our hands (data not shown). Seed plants may thus be too divergent from ferns to test Azolla TF-function.

Most Azolla phenolics had a distinct spatial distribution (**Figure 4 and 5**), moreover, transcripts from the different homologs encoding PBP/FBP-enzyme accumulated differently during cold-treatment (**File S2**). Flavonoid 3ʹ-hydroxylase (F3’H) controls the ratio of apigenidin and luteolinidin in *S. bicolor* (Mizuno et al., 2014). F3’H from Azolla, therefore, could be expressed more in the epidermis, accumulating mostly 3’-hydroxylated luteolinidin, than in the parenchyma, accumulating mostly apigenidin (**Figure 1B**; **Figure 4A-C**). F3’H from Azolla is difficult to identify because it is from the CYP450 superfamily and part takes in several pathways: (epi)catechin, the building block for Azolla PA-biosynthesis (Güngör et al., 2021), is also 3’-hydroxylated by an F3’H, and its accumulation resembled the pattern of apigenidin (**Figure 5B**). Different F3’H-homologs, therefore, likely control 3’-hydroxylation of DAs and PAs, which could further be segregated by compartmentalization in the cell (Brillouet et al., 2013).

To regulate PA-production (Yu et al., 2022), *Medicago truncatula*, *A. thaliana,* and *V. vinifera* have each recruited LAR, ANS and anthocyanidin reductase (ANR) differently. Predicting *in planta* function of Azolla FBP-enzymes based on seed plant data is therefore challenging. Moreover, Azolla may produce PAs yet by another pathway. To genetically engineer Azolla with less PAs for improved protein extractability, knocking-out TFs regulating the PBP, or pathways thereafter may be an easier approach. Phenolics accumulate at high concentrations even in green non-stressed Azolla, MYBs regulating the PBP and FBP may therefore be constitutively expressed. Overall, type VIII-D MYBs were the most expressed in Azolla and some were also upregulated in cold (**File S2**). Afi_v2_s2495G000440.1 peaked in **C14**, coinciding with the first redness in shoot apices, whilst Afi_v2_s25G003460.1 peaked in **C21**, when the whole fronds were bright red (**Table 2; Figure S7**). Afi_v2_s42G000270.4 had a similar expression in **C14** and **C21**, comparable to that of the differentially induced bHLH TF Afi_v2_s29G001620.1 (**Table 2, File S2**). Azolla type VIII-D MYBs, therefore, may play roles in PBP and FBP regulation.

L-type *Af*DFR1 had a substrate-interaction region similar to *Sb*FNR (**Figure 6B**), converted naringenin and eriodictyol *in vitro* characteristic for FNR-activity (**Figure 6C-D**); consistently, *Af*DFR1 did not substitute *At*DFR *in planta* which converts dihydroquercetin. *Af*DFR1 regulates DA-production in Azolla with a transcriptional off/on-switch in cold (**Figure 6A**). Q-type *De*DFR2 substituted *At*DFR and rescued PA-production (Chen et al., 2020); Q-type *Af*DFR2 could therefore catalyze PA-synthesis. *Af*DFR2 (Afi_v2_s5G012490.4) was, however, downregulated in cold (**Figure 6A**), whilst *Af*LAR was upregulated (**Table 2**). Reduced LAR expression correlated with less soluble PAs (Cannavò et al., 2023). Thus, when Azolla stops growing in cold, increased *Af*LAR may alter the degree of polymerization of existing PA-subunits (Liu et al., 2016), instead of channeling more substrate via *Af*DFR2.

Transcripts encoding GELP Afi_v2_s29G003930.6 and CYP450 Afi_v2_s35G001880.1 accumulated most highly in cold (**Table 2**), but do not encode the hypothesized second and final step of DA-biosynthesis. GELPs have been found to participate in the PBP, but are not well studied (Teutschbein et al., 2010; Miguel et al., 2020). Azolla GELPs could help producing different CQAs in cold. CYP450s are the biggest plant enzyme family and have their own nomenclature (Hansen et al., 2021); Afi_v2_s35G001880.1 was automatically annotated by Mercator as CYP736A12. Overexpression of a *Panax ginseng* homolog in *A. thaliana* increased tolerance to chlorotoluron, a phenolic herbicide (Khanom et al., 2019).

ANS has been considered for the final step in DA-biosynthesis. ANS is involved in both PA– and HA-biosynthesis in seed plants, as anthocyanidins are also a substrate for ANR (Dixon et al., 2005). *A. thaliana* produces PAs with only ANS and ANR, whilst *M. truncatula* and *V. vinifera* employ both ANS, ANR and LAR (Yu et al., 2022). Thus far, only LAR has been characterized in Azolla (Güngör et al., 2021), and more enzymes, such as ANS and ANR, are yet to be identified to propose a working model for Azolla PA-production and polymerization. ANR is highly similar to TKPR (Grienenberger *et al*. 2011). TKPR Afi_v2_s5G008050.13 had a higher transcript accumulation than all *Af*DFRs and *Af*LAR in all samples, and was upregulated in cold, following a similar trend as *Af*LAR (**Figure 6A**, **Table 2, File S2**). TKPR, therefore, possibly participates in PA-production.

ANS belongs to the 2OGD-superfamily together with FBP-enzymes FLS, flavone synthase (FNS) and flavanone 3-hydroxylase (F3H) (Kawai et al., 2014). FNS and F3H, respectively, use flavanones to produce flavones detected in Azolla roots (Teixeira et al., 2001), and dihydroflavonols (Yonekura-Sakakibara et al., 2019). Bifunctional FNS/F3H were characterized in bryophytes and lycophytes and likely form the evolutionary basis for 2OGD-type FBP-enzymes (Li et al., 2020). Azolla therefore likely possesses these enzymes as well, especially since dihydroflavonols are also the substrate for DFR towards PA-production. FLS uses dihydroflavonols to produce flavonols such as kaempferol and quercetin, which have been detected in Azolla (Güngör et al., 2021). None of the seven FBP-related Azolla 2OGD-enzymes were responsive to cold to the extent of *Af*DFR1 (**Figure 6A**, **Figure 7B**). The enzyme that performs the final step in DA-production might thus already be expressed in green ferns with *Af*DFR1 redirecting the substrates upon stress. Non-enzymatic oxidation of the colorless precursor is also a possibility, as reported for some HAs (Yoshida et al., 2020). To uncover all enzymes of DA-biosynthesis spatially resolved transcript-profiling or an untargeted approach will provide more insight.

## Acknowledgments

We thank Dr. Cristina Sarasa-Buisan for her help interpreting *N. azollae* gene-expression, Kelvin Adema for his help isolating leaf pockets, Javier Sastre Toraño for the LC-MS measurements of the *in vitro* enzyme assays and Laura W. Dijkhuizen for the phylogenetic analysis of the 2OGD-superfamily.

## Funding

We acknowledge funding for EG from the NWO-TTW grant AZOPRO (project number 16294). The LDI-MSI has been made possible with the support of the Dutch Province of Limburg and a Netherlands Organization for Scientific Research VIDI grant (project number TTW VI.Vidi.198.011) awarded to Shane R. Ellis.

## Author contributions

EG grew the plants, performed the physiological experiments, and followed the development of the red color and *N. azollae* during cold-treatment. EG performed the dual RNA-Seq, with help of HS and GB, and cloned and assayed *Af*DFR1. BB performed the LDI-MSI on plants provided by EG, in consultation with RH and SE. EG wrote the manuscript together with HS and with input from all other authors. All authors agreed to the list of authors and identified contributions from each other.

## Notes

### Competing Interest Statement

The authors have declared no competing interest.

