## Supplementary Materials for "High-resolution metabolite imaging: luteolinidin accumulates at the host-cyanobiont interface during cold-acclimation in *Azolla* symbioses"

**Supplementary Figures**

**Figure S1** Recovery of red *A. filiculoides* after cold treatment (60-days at 4°C) over a period of 5 weeks at 21°C.

**Figure S2** Typical growth habit of different Azolla species during 21 days 4°C cold-treatment on demineralized water ($H_2O$), SM, SM with 5 µM ABA and SM without phosphate (-P), and during 14 days of recovery at 21°C on SM.

**Figure S3** Typical phenotype of *A. filiculoides* and *N. azollae* during 21 days of cold-treatment.

**Figure S4** Typical phenotype of *A. filiculoides* and *N. azollae* during 21-days of recovery at 21°C, after 21-days 4°C of cold-treatment as per **Figure S3**.

**Figure S5** DMACA-staining for proanthocyanidins in the *A. filiculoides* sporophyte.

**Figure S6** High-resolution overview of the ROI used for LDI-MSI acquisition in **Figures 4-5**.

**Figure S7** Phenotypes of the cold-acclimated *A. filiculoides* symbioses used for dual RNA-Seq profiling.

**Figure S8** Uniquely mapped reads on the *N. azollae* and *A. filiculoides* genomes from the dual RNA-Seq experiment from **Figure S7**.

**Figure S9** Expression pattern in log2 Fold Change of the 237 differentially expressed *N. azollae* genes after cold treatment and after development of the reproductive stages.

**Supplementary Files (submitted separately, .xlsx)**

**File S1** Transcript accumulation in RPM and DESeq2 analysis for *N. azollae* genes during cold-treatment and in the reproductive structures.

**File S2** Transcript accumulation in RPM and DESeq2 analysis for *A. filiculoides* genes defined by Afi_v2 during cold-treatment.

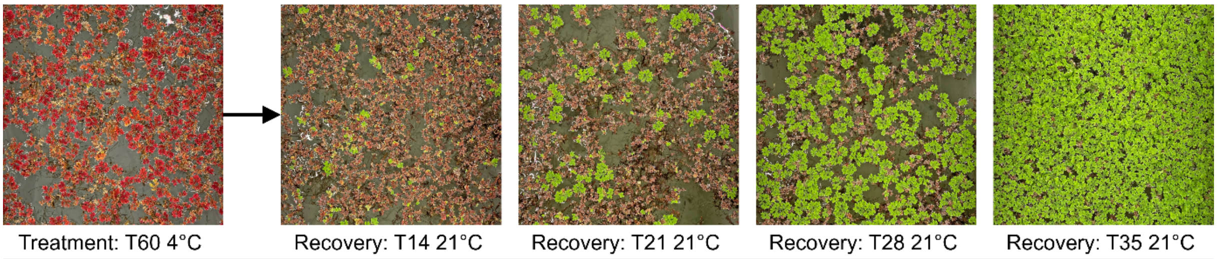


**Figure S1** Recovery of red *A. filiculoides* after cold treatment (60-days at 4°C) over a period of 5 weeks at 21°C, continuously on the same standard medium.

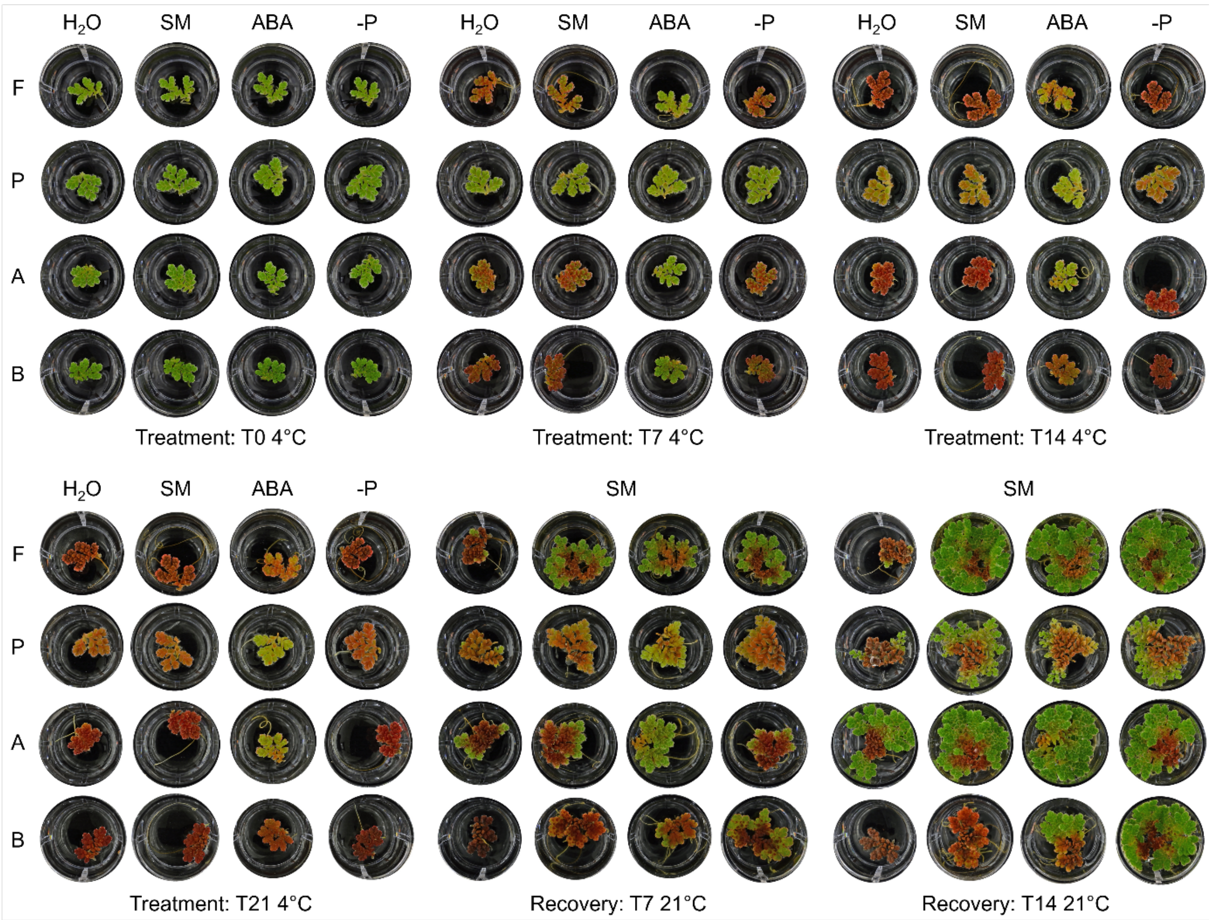


**Figure S2** Typical growth habit of different Azolla species during 21 days 4°C cold-treatment on demineralized water ($H_2O$), SM, SM with 5 µM ABA and SM without phosphate (-P), and during 14 days of recovery at 21°C on SM. Images are representative from three independent biological repeats and were taken after 0 (T0), 7 (T7), 14 (T14), and 21 (T21) days in the cold at 4°C, followed by 7 (T7) and 14 (T14) days at 21°C. Species, F: *A. filiculoides*, P: *A. pinnata*, A: *Azolla sp. Anzali*, B: *Azolla sp. Bordeaux*.

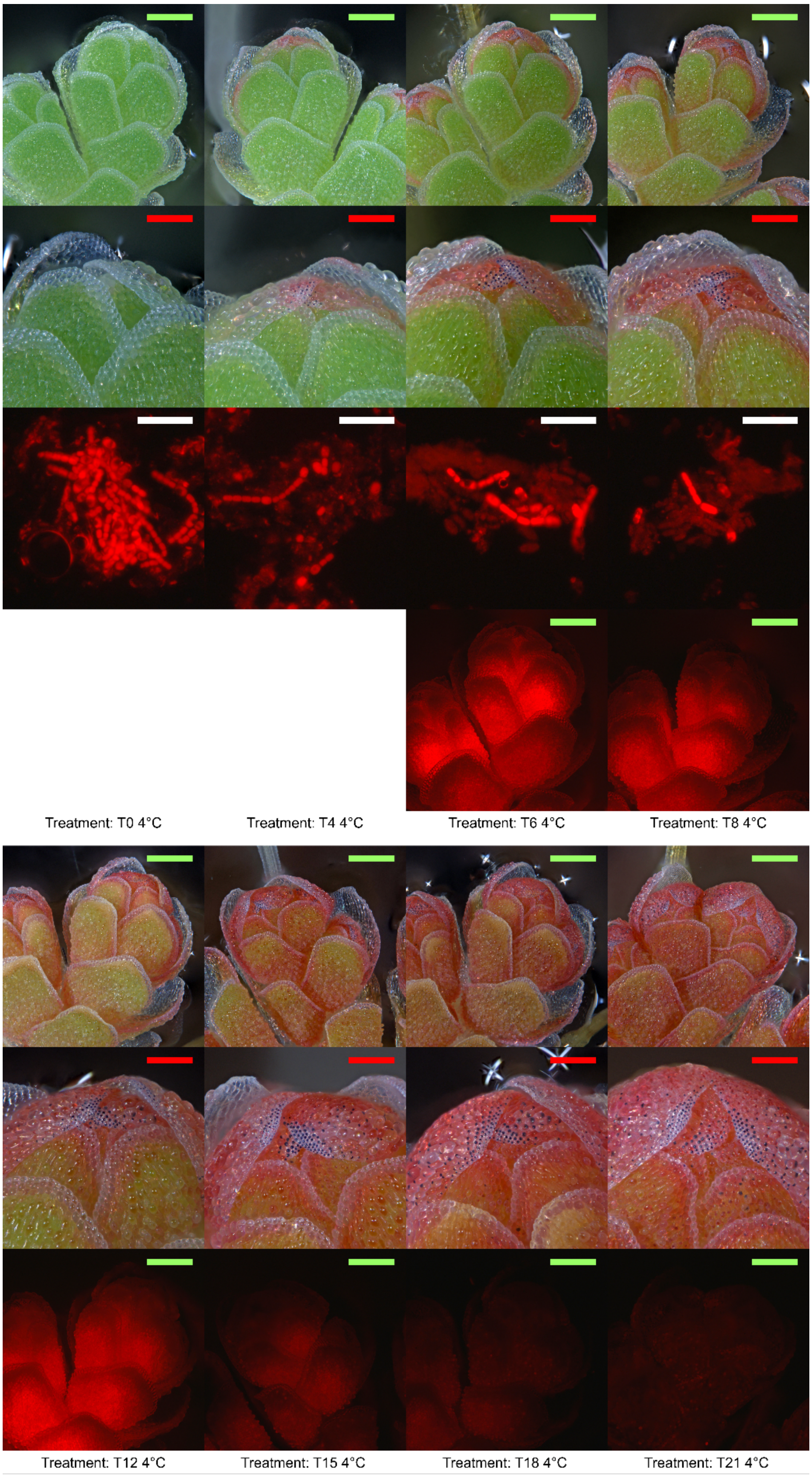


**Figure S3** Typical phenotype of *A. filiculoides* and *N. azollae* during 21 days of cold-treatment. From top to bottom: branch, close-up shoot apex, cyanobacterial filaments (if detectable) and otherwise branch under RFP-settings. Scale bars: green, red and white, respectively, correspond to 500 µm, 200 µm and 100 µm.

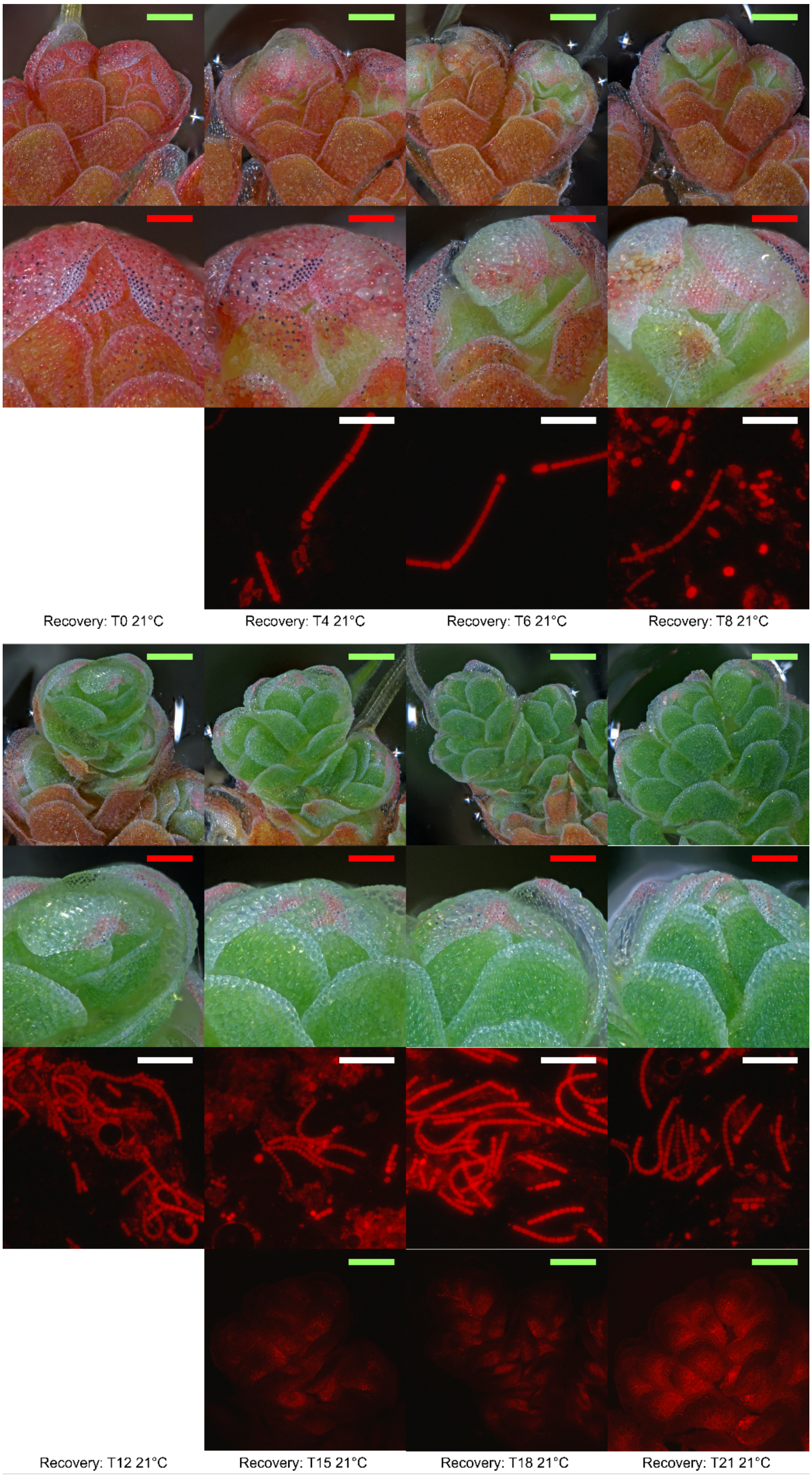


**Figure S4** Typical phenotype of *A. filiculoides* and *N. azollae* during 21-days of recovery at 21°C, after 21-days 4°C of cold-treatment as per **Figure S3**. From top to bottom: branch, close-up shoot apex, cyanobacterial filaments (if detectable) and branch under RFP-settings. Scale bars: green, red and white, respectively, correspond to 500 µm, 200 µm and 100 µm.

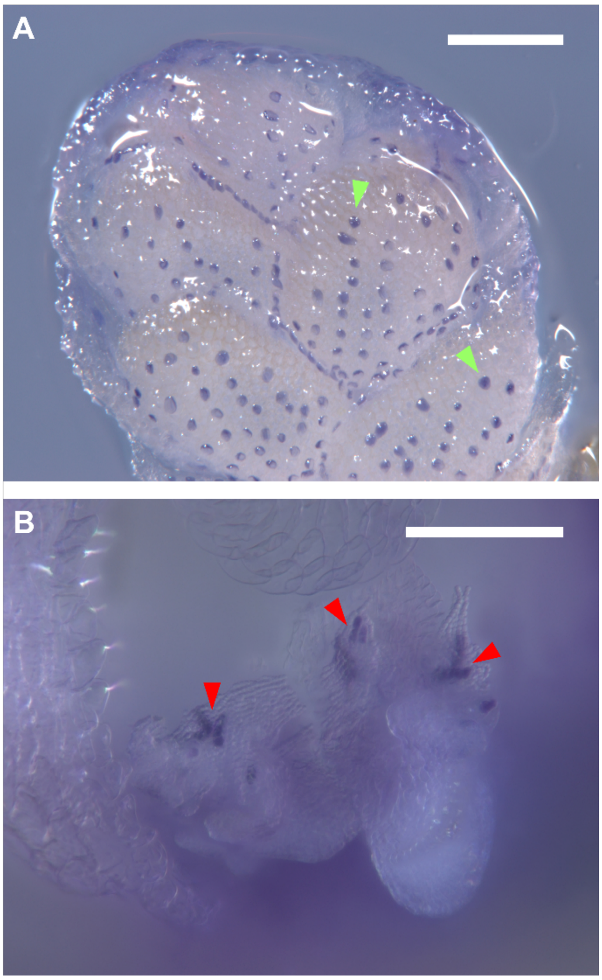


**Figure S5** DMACA-staining for proanthocyanidins in the *A. filiculoides* sporophyte. **A** Branch with the translucent papillae (green arrows) and hyaline border cells at the leaf rim turning especially purple. **B** Exposed leaf pocket trichomes (red arrows) stained purple and were surrounded by translucent cyanobacterial filaments. Scale bars correspond to 300 µm.

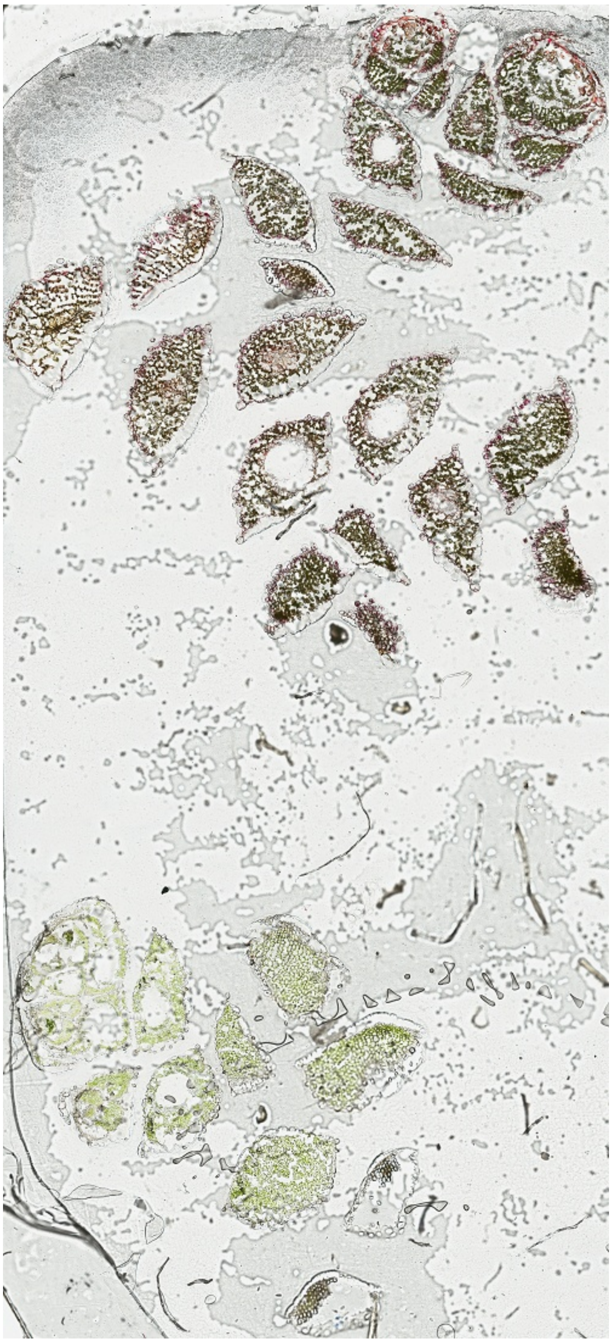

**Figure S6** High-resolution image of the ROI used for LDI-MSI acquisition in **Figures 4-5**.

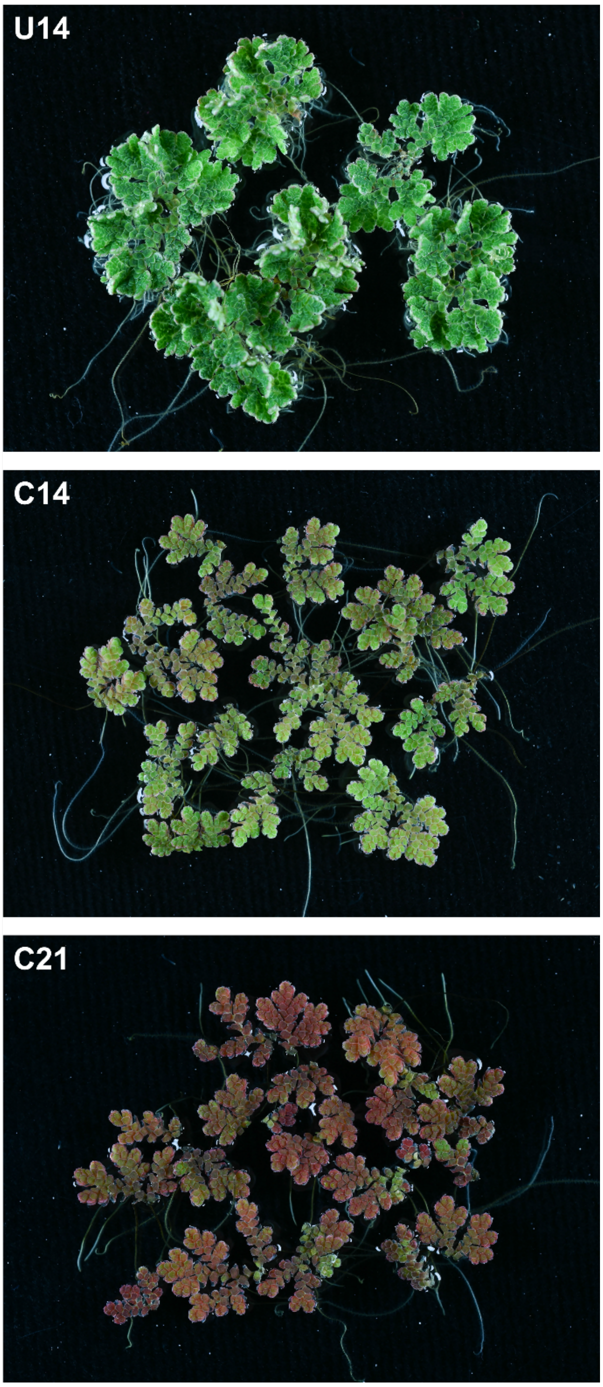


**Figure S7** Phenotypes of the cold-acclimated *A. filiculoides* symbioses used for dual RNA-Seq profiling. From top to bottom: untreated (**U14**), 14-days cold-treated (**C14**) and 21-days cold-treated (**U21**).

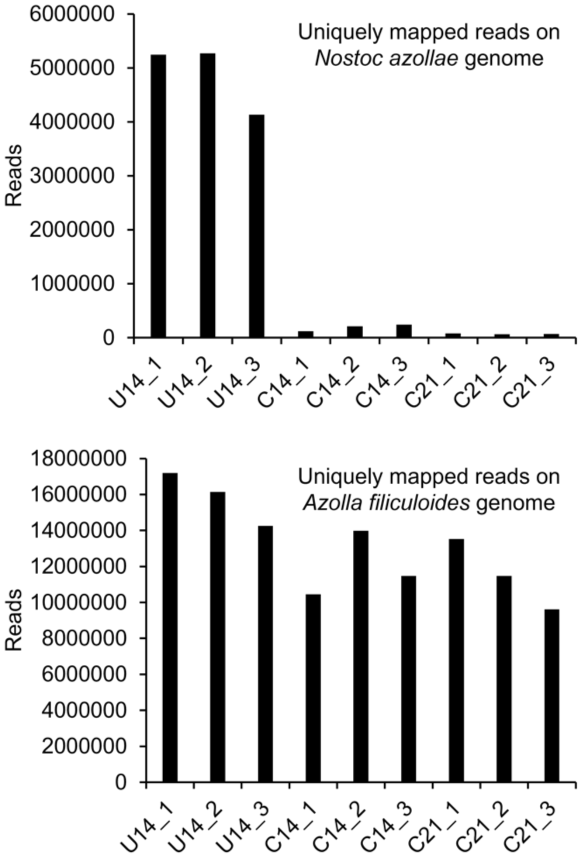


**Figure S8** Uniquely mapped reads on the *N. azollae* and *A. filiculoides* genomes, from the dual RNA-Seq experiment from **Figure S7**.

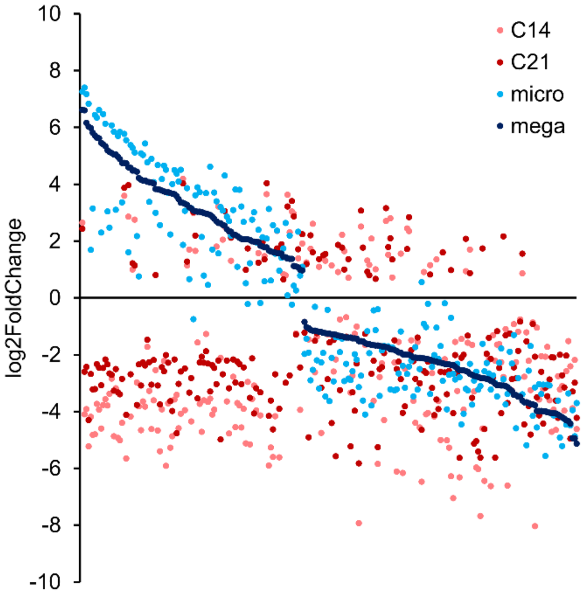


**Figure S9** Expression pattern in log2 Fold Change of 237 differentially expressed *N. azollae* genes (padj<0.1) after cold treatment (**C14** and **C21**, compared to **U14**) and after development of the reproductive stages (**mega**sporocarps and **microcro**sporocarps, compared to **sp**orophytes; Güngör et al., 2023). The log2 Fold Change was sorted from highest to lowest for sample **mega**. Source data in **File S1**.

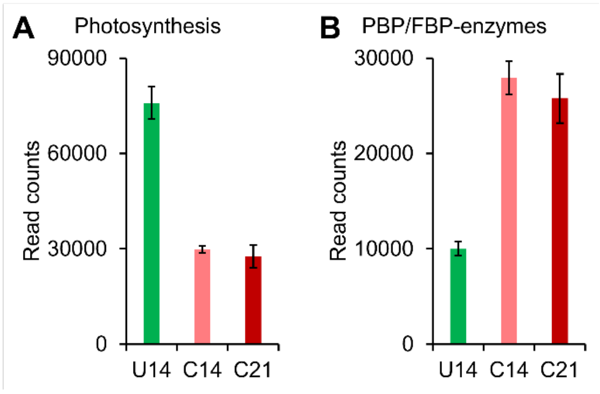


**Figure S10** Total transcript investment of *A. filiculoides* during cold-treatment (untreated, **U14**; 14-days cold-treated, **C14**; and 21-days cold-treated, **U21**) for 239 genes involved in photosynthesis as determined by annotation with Mercator (Lohse et al., 2013), and 60 manually-curated genes involved in the PBP and FBP. Source data in **File S2**.

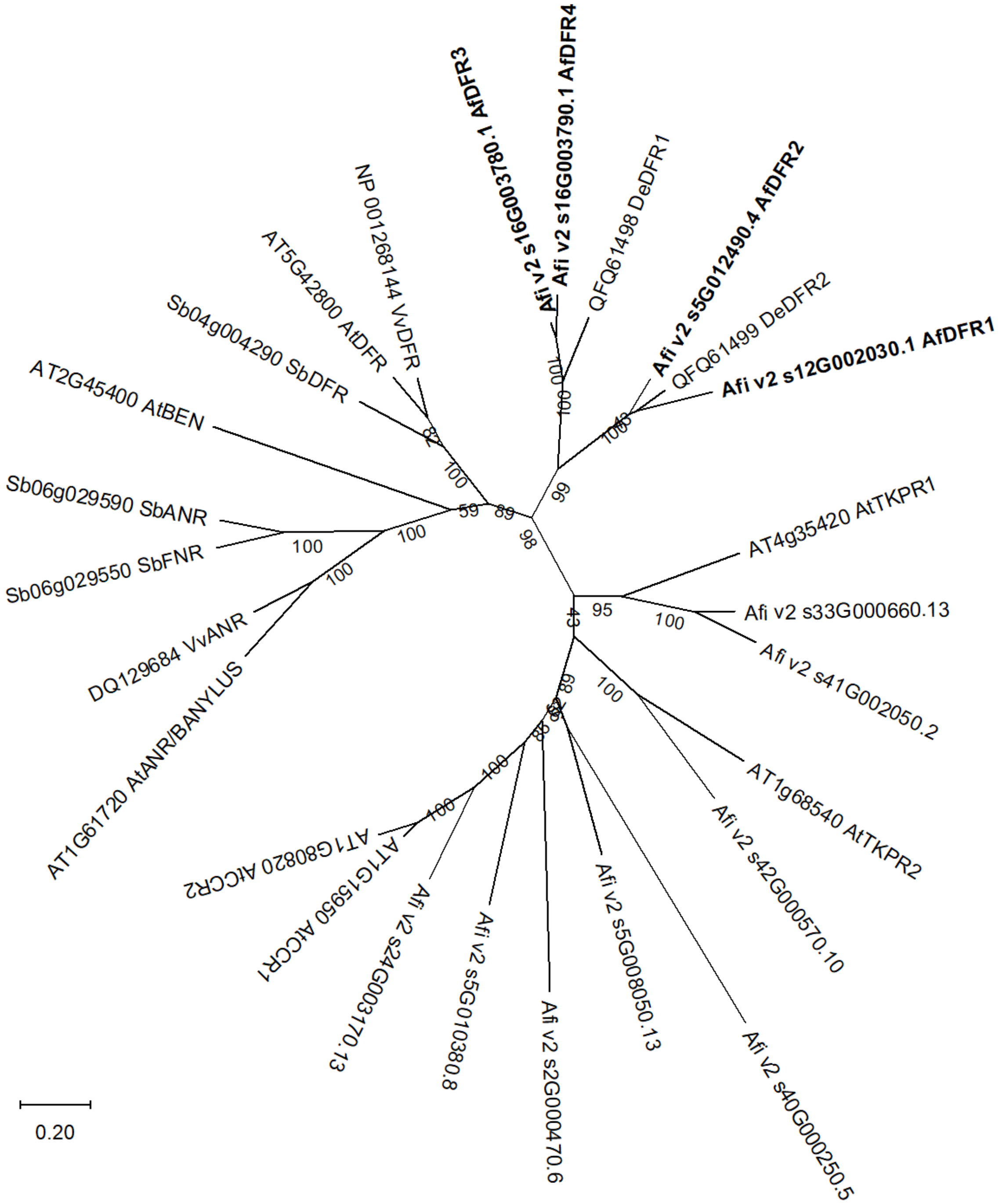


**Figure S11** Phylogenetic analysis of 13 DFR-related *A. filiculoides* genes found in the Afi_v2 genome. Functionally characterized angiosperm and fern sequences were from Grienenberger et al., 2011, Chen et al., 2020 and Lewis et al., 2023. Accession numbers or aliases of the sequences are provided in the tree. MEGA11 was used for both the alignment with MUSCLE and the generation of a maximum-likelihood tree, values for branch support were obtained from 100 times bootstrapping (Tamura et al., 2021). The scale corresponds to 0.2 average substitutions per site.

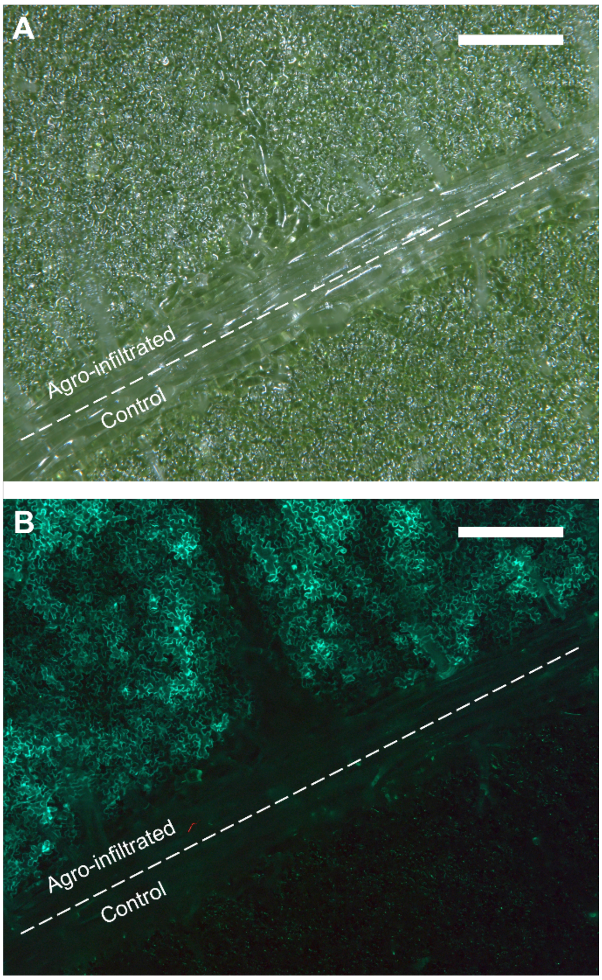


**Figure S12** Typical GFP-expression in *N. tabacum* leaves 72 hours after Agro-infiltration with the 35S::*Af*DFR1 plasmid. **A** Leaf under brightfield microscopy and **B** under Zeiss GFP filter set 38HE (excitation 470 nm, emission 525 nm). The diagonal vein separates a part of the leaf that was kept untouched as control. Scale bars correspond to 400 µm.
